# Genomic diversity affects the accuracy of bacterial SNP calling pipelines

**DOI:** 10.1101/653774

**Authors:** Stephen J. Bush, Dona Foster, David W. Eyre, Emily L. Clark, Nicola De Maio, Liam P. Shaw, Nicole Stoesser, Tim E. A. Peto, Derrick W. Crook, A. Sarah Walker

## Abstract

**Background:** Accurately identifying SNPs from bacterial sequencing data is an essential requirement for using genomics to track transmission and predict important phenotypes such as antimicrobial resistance. However, most previous performance evaluations of SNP calling have been restricted to eukaryotic (human) data. Additionally, bacterial SNP calling requires choosing an appropriate reference genome to align reads to, which, together with the bioinformatic pipeline, affects the accuracy and completeness of a set of SNP calls obtained.

This study evaluates the performance of 41 SNP calling pipelines using simulated data from 254 strains of 10 clinically common bacteria and real data from environmentally-sourced and genomically diverse isolates within the genera *Citrobacter, Enterobacter, Escherichia* and *Klebsiella*.

**Results:** We evaluated the performance of 41 SNP calling pipelines, aligning reads to genomes of the same or a divergent strain. Irrespective of pipeline, a principal determinant of reliable SNP calling was reference genome selection. Across multiple taxa, there was a strong inverse relationship between pipeline sensitivity and precision, and the Mash distance (a proxy for average nucleotide divergence) between reads and reference genome. The effect was especially pronounced for diverse, recombinogenic, bacteria such as *Escherichia coli*, but less dominant for clonal species such as *Mycobacterium tuberculosis*.

**Conclusions:** The accuracy of SNP calling for a given species is compromised by increasing intra-species diversity. When reads were aligned to the same genome from which they were sequenced, among the highest performing pipelines was Novoalign/GATK. However, across the full range of (divergent) genomes, among the consistently highest-performing pipelines was Snippy.

## Introduction

Accurately identifying single nucleotide polymorphism (SNPs) from bacterial DNA is essential for monitoring outbreaks (as in [1, 2]) and predicting phenotypes, such as antimicrobial resistance [3], although the pipeline selected for this task strongly impacts the outcome [4]. Current bacterial sequencing technologies generate short fragments of DNA sequence (‘reads’) from which the bacterial genome can be reconstructed. Reference-based mapping approaches use a known reference genome to guide this process, using a combination of an aligner, which identifies the location in the genome each read is likely to have arisen from, and a variant caller, which summarises the available information at each site to identify variants including SNPs and indels (see reviews for an overview of alignment [5, 6] and SNP calling [7] algorithms). This evaluation focuses only on SNP calling; we did not evaluate indel calling as this can require different algorithms (see review [8]). The output from different aligner/caller combinations is often poorly concordant. For example, up to 5% of SNPs are uniquely called by one of five different pipelines [9] with even lower agreement upon structural variants [10].

Although a mature field, systematic evaluations of variant calling pipelines are often limited to eukaryotic data, usually human [11–15] but also *C. elegans* [16] and dairy cattle [17] (see also review [18]). This is because truth sets of known variants, such as the Illumina Platinum Genomes [19], are relatively few in number and human-centred, being expensive to create and biased toward the methods that produced them [20]. As such, to date, bacterial SNP calling evaluations are comparatively limited in scope (for example, comparing 4 aligners with 1 caller, mpileup [21], using *Listeria monocytogenes* [22]).

Relatively few truth sets exist for bacteria and so the choice of pipeline for bacterial SNP calling is often informed by performance on human data. Many evaluations conclude in favour of the publicly-available BWA-mem [23] or commercial Novoalign (www.novocraft.com) as choices of aligner, and GATK [24, 25] or mpileup as variant callers, with recommendations for a default choice of pipeline, independent of specific analytic requirements, including Novoalign followed by GATK [26], and BWA-mem followed by either mpileup [14], GATK [12], or VarDict [11].

This study evaluates a range of SNP calling pipelines across multiple bacterial species, both when reads are sequenced from and aligned to the same genome, and when reads are aligned to a representative genome of that species. In order to cover a broad range of methodological approaches, we assessed the combination of 4 commonly used short read aligners (BWA-mem [23], minimap2 [27], Novoalign and Stampy [28]) and 10 variant callers (16GT [29], Freebayes [30], GATK HaplotypeCaller [24, 25], LoFreq [31], mpileup [21], Platypus [32], SNVer [33], SNVSniffer [34], Strelka [35] and VarScan [36]), alongside Snippy (https://github.com/tseemann/snippy), a haploid core variant calling pipeline constituting a bespoke aligner/caller combination of BWA-mem, minimap2, and Freebayes. Reasons for excluding other programs are detailed in Supplementary Text 1.

To evaluate each pipeline, we simulated 3 sets of 150bp and 3 sets of 300bp reads (characteristic of the Illumina NextSeq and MiSeq platforms, respectively) at 50-fold depth from 254 strains of 10 clinically common species (2 to 36 strains per species), each with fully sequenced (closed) core genomes: the Gram-positive *Clostridioides difficile* (formerly *Clostridium difficile* [37]), *Listeria monocytogenes, Staphylococcus aureus*, and *Streptococcus pneumoniae* (all Gram-positive), *Escherichia coli, Klebsiella pneumoniae, Neisseria gonorrhoeae, Salmonella enterica*, and *Shigella dysenteriae* (all Gram-negative), and *Mycobacterium tuberculosis*. For each strain, we evaluated all pipelines using two different genomes for alignment: one being the same genome from which the reads were simulated, and one being the NCBI ‘reference genome’, a high-quality (but essentially arbitrary) representative of that species, typically chosen on the basis of assembly and annotation quality, available experimental support, and/or wide recognition as a community standard (such as *C. difficile* 630, the first sequenced strain for that species [38]). We added approximately 8000-25,000 SNPs *in silico* to each genome, equivalent to 5 SNPs per genic region, or 1 SNP per 60-120 bases.

While simulation studies can offer useful insight, they can be sensitive to the specific details of the simulations. Therefore, we also evaluated performance on real data to verify our conclusions. We used 16 environmentally-sourced and genomically diverse Gram-negative species of the genera *Citrobacter, Enterobacter, Escherichia* and *Klebsiella*, along with two reference strains, from which closed hybrid *de novo* assemblies were previously generated using both Illumina (short) and ONT (long; Oxford Nanopore Technologies) reads [39].

All pipelines aim to call variants with high specificity (i.e. high proportion of non-variant sites in the truth set correctly identified as the reference allele by the pipeline) and high sensitivity (i.e. high proportion of true SNPs found by the pipeline, a.k.a. recall). The optimal trade-off between these two properties may vary depending on the application. For example, in transmission inference, minimising false positive SNP calls (i.e. high specificity), is likely to be most important, whereas high sensitivity may be more important when identifying variants associated with antibiotic resistance. We therefore report detailed performance metrics for all pipelines, including recall/sensitivity, precision (a.k.a. positive predictive value, the proportion of SNPs identified that are true SNPs), and the F-score, the harmonic mean of precision and recall [40].

## Results

### Evaluating SNP calling pipelines when the genome for alignment is also the source of the reads

The performance of 41 SNP calling pipelines (Supplementary Table 1) was first evaluated using reads simulated from 254 closed bacterial genomes (Supplementary Table 2), as illustrated in Figure 1. In order to exclude biases introduced during other parts of the workflow, such as DNA library preparation and sequencing error, reads were simulated error-free. There was negligible difference in performance when reads were simulated with sequencing errors (see Supplementary Text 1).

**Figure 1.**
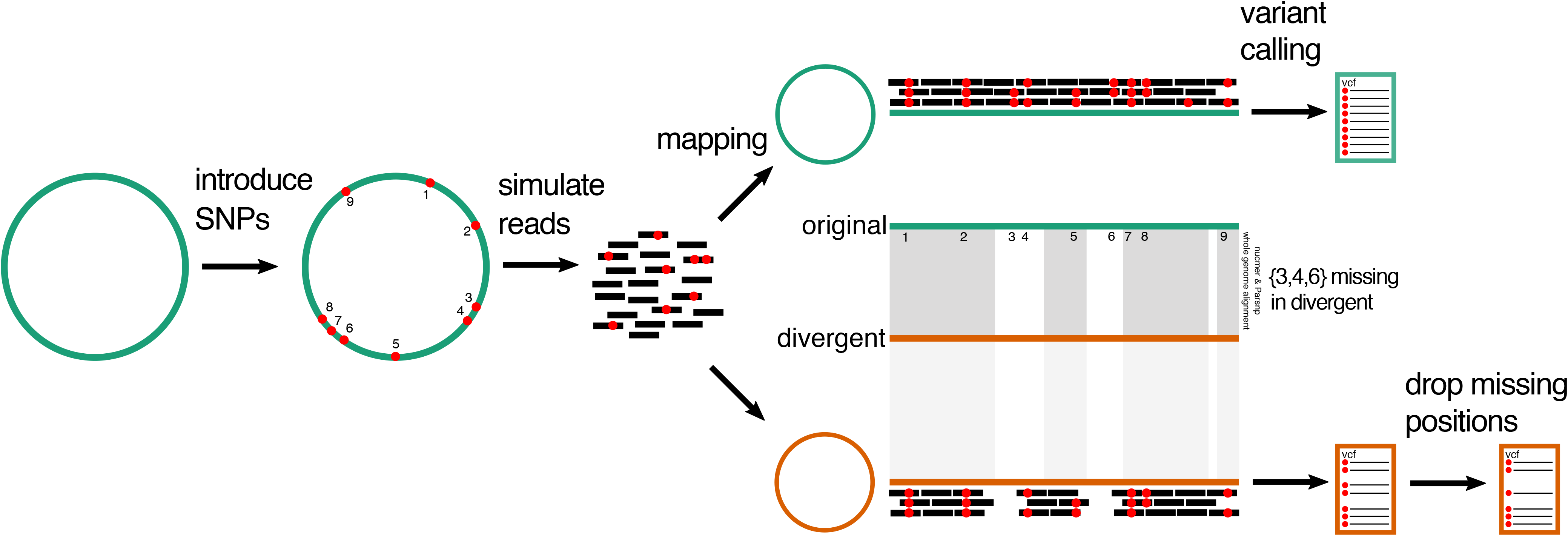
Overview of SNP calling evaluation. SNPs were introduced *in silico* into 254 closed bacterial genomes (Supplementary Table 2) using Simulome. Reads were then simulated from these genomes. 41 SNP calling pipelines (Supplementary Table 1) were evaluated using two different genomes for read alignment: the original genome from which the reads were simulated and a divergent genome, the species-representative NCBI ‘reference genome’. In the latter case, it will not be possible to recover all of the original *in silico* SNPs as some will be found only within genes unique to the original genome. Accordingly, to evaluate SNP calls, the coordinates of the original genome need to be converted to those of the representative genome. To do so, whole genome alignments were made using both nucmer and Parsnp, with consensus calls identified within one-to-one alignment blocks. Inter-strain SNPs (those not introduced *in silico*) are excluded. The remaining subset of *in silico* calls comprise the truth set for evaluation. There is a strong correlation between the total number of SNPs introduced *in silico* into the original genome and the total number of nucmer/Parsnp consensus SNPs in the divergent genome (Supplementary Figure 3).

This dataset contains 62,484 VCFs (comprising 2 read lengths [150 and 300bp] * 3 replicates * 254 genomes * 41 pipelines). The number of reads simulated from each species and the performance statistics for each pipeline – the number of true positives (TP), false positives (FP) and false negatives (FN), precision, recall, F-score, and total number of errors (i.e. FP + FN) per million sequenced bases – are given in Supplementary Table 3, with the distribution of F-scores illustrated in Figure 2A.

**Figure 2.**
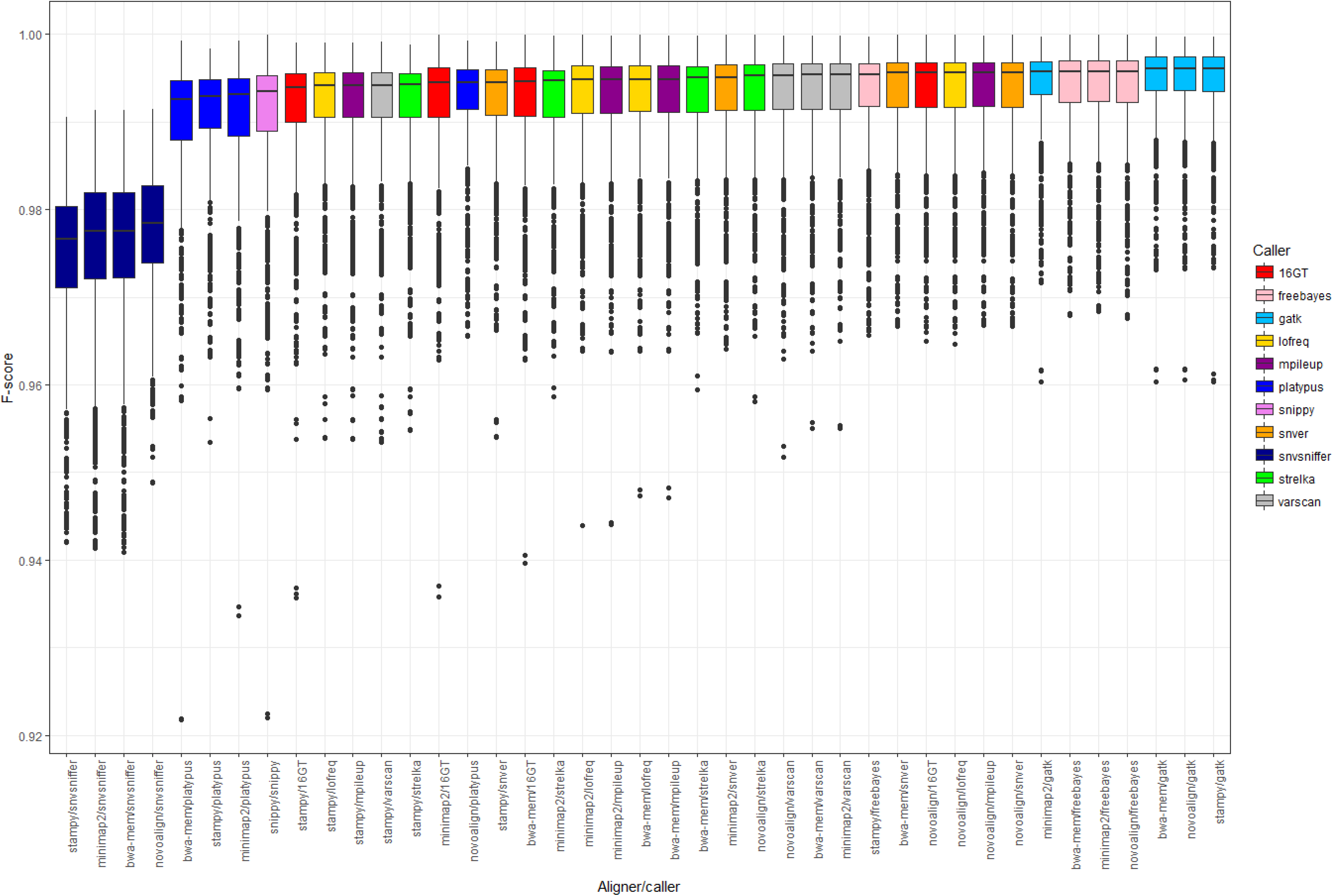
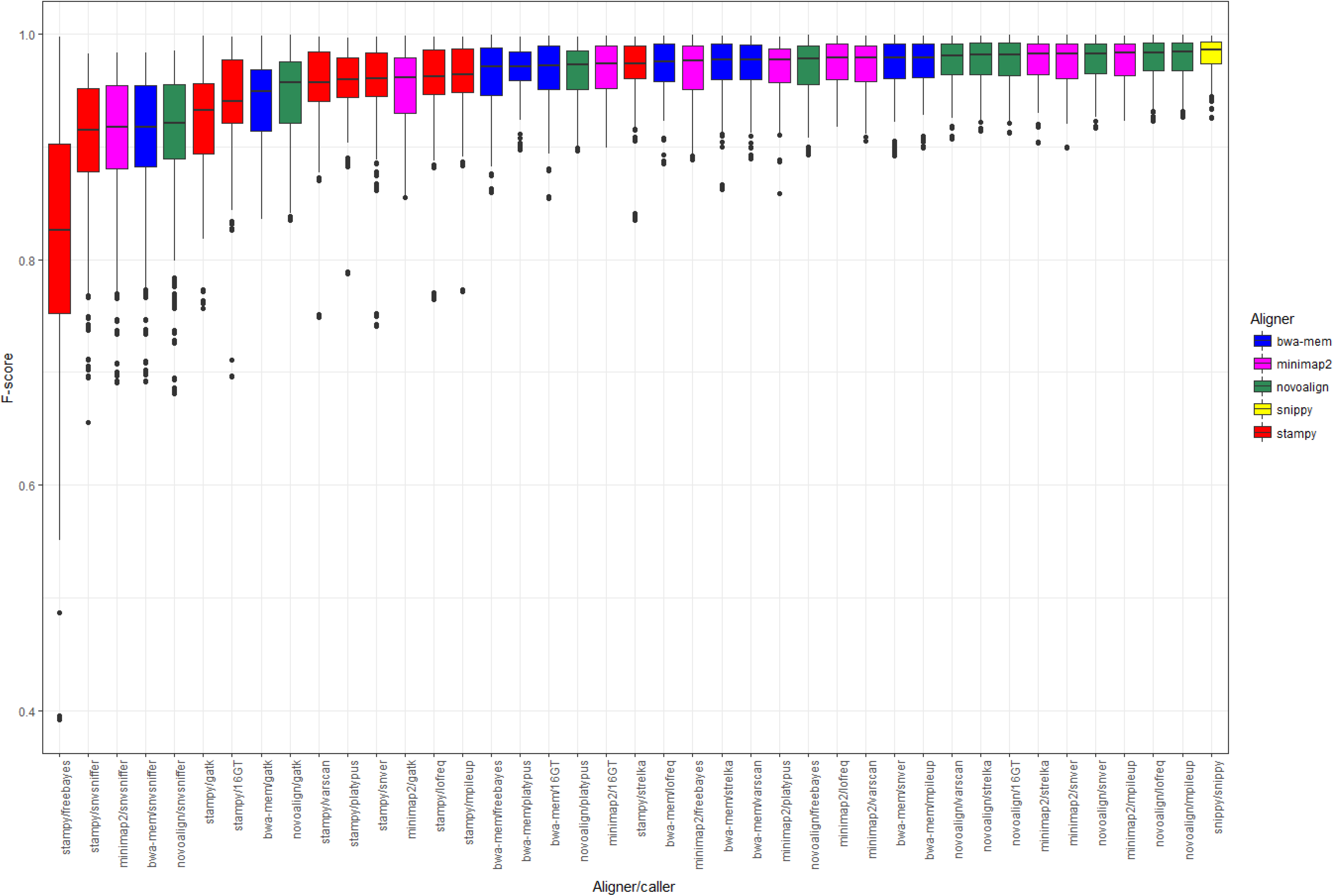
Median F-score per pipeline when the reference genome for alignment is (A) the same as the source of the reads, and (B) a representative genome for that species. Panels show the median F-score of 41 different pipelines when SNPs are called using error-free 150bp and 300bp reads simulated from 254 genomes (of 10 species) at 50-fold coverage. Pipelines are ordered according to median F-score and coloured according to either the variant caller (A) or aligner (B) in each pipeline. Note that because F-scores are uniformly > 0.9 when the reference genome for alignment is the same as the source of the reads, the vertical axes on each panel have different scales. Genomes are detailed in Supplementary Table 2, summary statistics for each pipeline in Supplementary Tables 3 and 6, and performance ranks in Supplementary Tables 4 and 7, for alignments to the same or to a representative genome, respectively.

Median F-scores were over 0.99 for all but four aligner/callers with small interquartile ranges (approx. 0.005), although outliers were nevertheless notable (Figure 2A), suggesting that reference genome can affect performance of a given pipeline.

Table 1 shows the top ranked pipelines averaged across all species’ genomes, based on 7 different performance measures and on the sum of their ranks (which constitutes an ‘overall performance’ measure, lower values indicating higher overall performance). Supplementary Table 4 shows the sum of ranks for each pipeline per species, with several variant callers consistently found among the highest-performing (Freebayes and GATK) and lowest-performing pipelines (16GT and SNVSniffer), irrespective of aligner.

**Table 1.** Summary of pipeline performance across all species’ genomes.

If considering performance across all species, Novoalign/GATK has the highest median F-score (0.994), lowest sum of ranks (10), the lowest number of errors per million sequenced bases (0.944), and the largest absolute number of true positive calls (15,778) (Table 1). However, in this initial simulation, as the reads are error-free and the reference genome is the same as the source of the reads, many pipelines avoid false positive calls and report a perfect precision of 1.

### Evaluating SNP calling pipelines when the genome for alignment diverges from the source of the reads

Due to the high genomic diversity of some bacterial species, the appropriate selection of reference genomes is non-trivial. To assess how pipeline performance is affected by divergence between the source and reference genomes, SNPs were re-called after mapping all reads to a single representative genome for that species (illustrated in Figure 1). To identify true variants, closed genomes were aligned against the representative genome using both nucmer [41] and Parsnp [42], with consensus calls identified within one-to-one alignment blocks (see Methods). Estimates of the distance between each genome and the representative genome are given in Supplementary Table 2, with the genomic diversity of each species summarised in Supplementary Table 5. We quantified genomic distances using the Mash distance, which reflects the proportion of k-mers shared between a pair of genomes as a proxy for average nucleotide divergence [43]. The performance statistics for each pipeline are shown in Supplementary Table 6, with an associated ranked summary in Supplementary Table 7.

In general, aligning reads from one strain to a divergent reference leads to a decrease in median F-score and increase in interquartile range of the F-score distribution, with pipeline performance more negatively affected by choice of aligner than caller (Figure 2B).

Although across the full range of genomes, many pipelines show comparable performance (Figure 2B), there was a strong negative correlation between the Mash distance and F-score (Spearman’s *rho* = −0.72, p < 10^−15^; Figure 3A). The negative correlation between F-score and the total number of SNPs between the strain and representative genome, i.e. the set of strain-specific *in silico* SNPs plus inter-strain SNPs, was slightly weaker (*rho* = −0.58, p < 10^−15^; Supplementary Figure 1). This overall reduction in performance with increased divergence was more strongly driven by reductions in recall (i.e., by an increased number of false negative calls) rather than precision as there was a particularly strong correlation between distance and recall (Spearman’s *rho* = −0.94, p < 10^−15^; Supplementary Figure 2).

**Figure 3.**
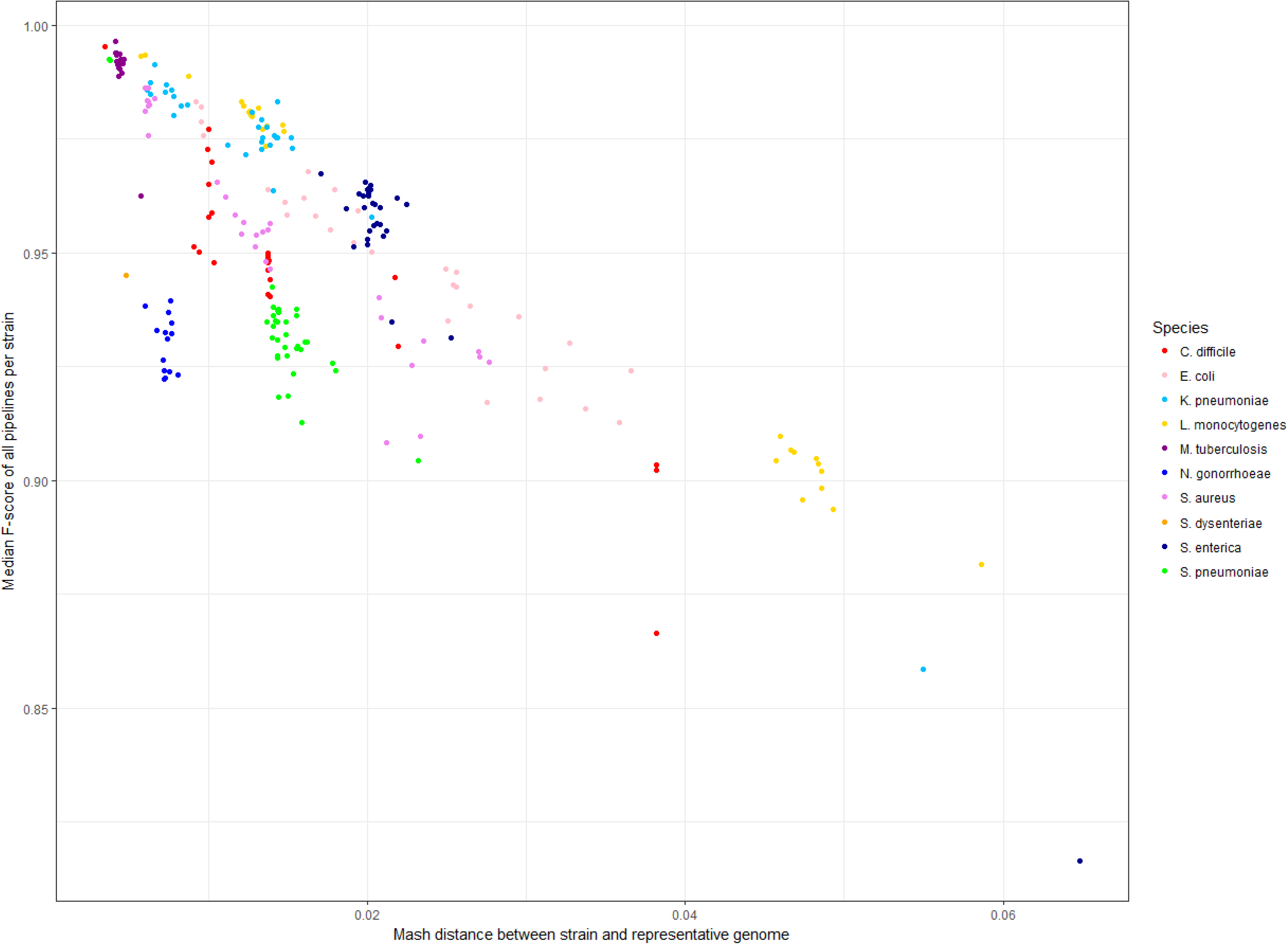
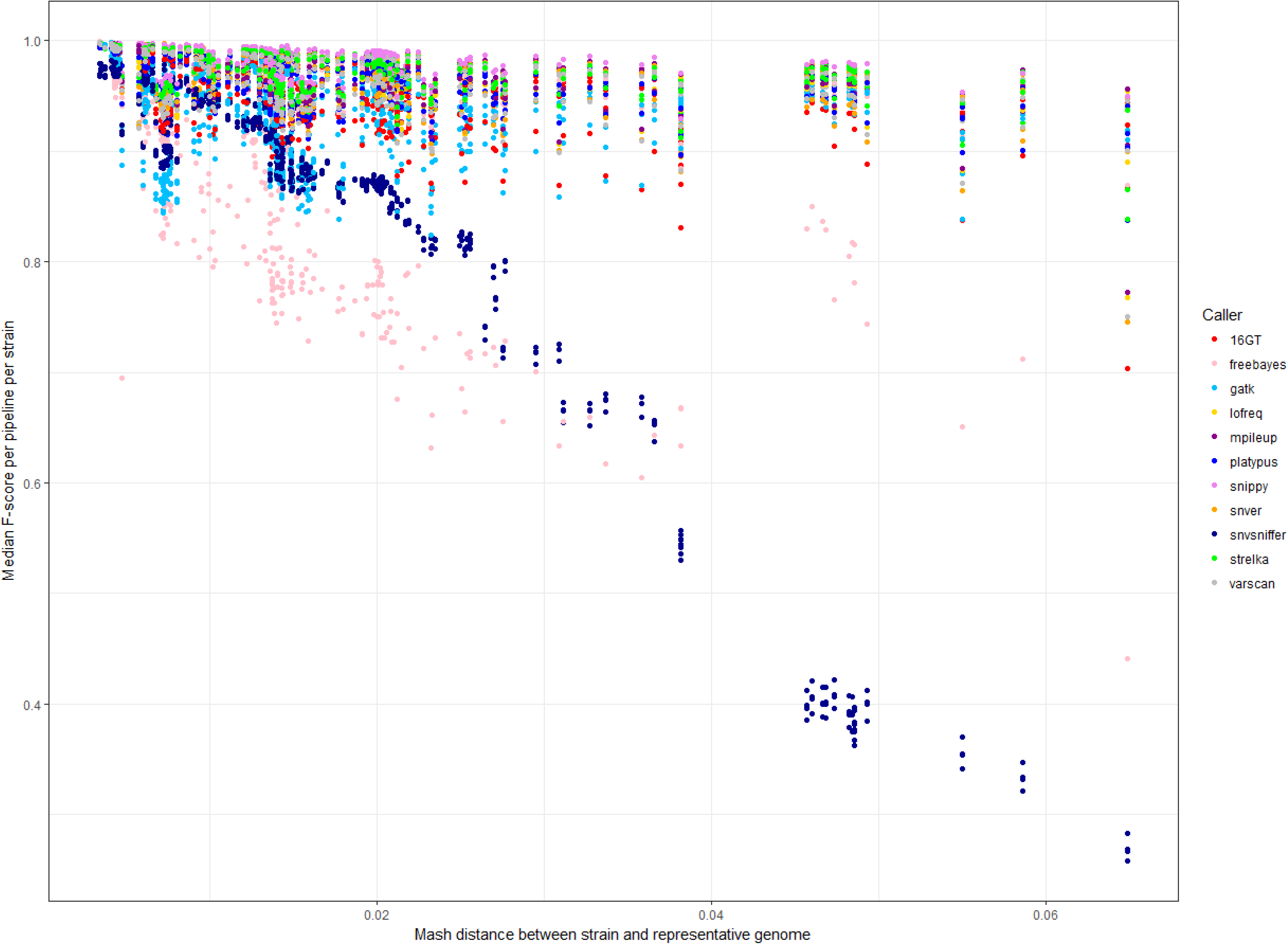
Reduced performance of SNP calling pipelines with increasing genetic distance between the reads and the reference genome. Panel A shows that the median F-score across the complete set of 41 pipelines, per strain, decreases as the distance between the strain and the reference genome increases (assayed as the Mash distance, which is based on the proportion of k-mers shared between genomes). Each point indicates the median F-score, across all pipelines, for the genome of one strain per species (n = 254 strains). Points are coloured by the species of each strain (n = 10 species). Panel B shows the median F-score per pipeline per strain, with points coloured according to the variant caller in each pipeline. This shows that the performance of some SNP calling pipelines is more negatively affected by increasing distance from the reference genome. Summary statistics for each pipeline are shown in Supplementary Table 6, performance ranks in Supplementary Table 7 and the genetic distance between strains in Supplementary Table 2. Quantitatively similar results are seen if assaying distance as the total number of SNPs between the strain and representative genome, i.e. the set of strain-specific *in silico* SNPs plus inter-strain SNPs (Supplementary Figure 1).

Three commonly used pipelines – BWA-mem/Freebayes, BWA-mem/GATK and Novoalign/GATK – were among the highest performers when the reference genome is also the source of the reads (Table 1 and Supplementary Table 4). However, when the reference diverges from the reads, then considering the two ‘overall performance’ measures across the set of 10 species, Snippy instead has both the lowest sum of ranks (20) and the highest median F-score (0.982), along with the lowest number of errors per million sequenced bases (2.6) (Table 1).

Performance per species is shown in Table 2, alongside both the overall sum and range of these ranks per pipeline. Pipelines featuring Novoalign were, in general, consistently high-performing across the majority of species (that is, having a lower sum of ranks), although were outperformed by Snippy, which had both strong and uniform performance across all species (Table 2). By contrast, pipelines with a larger range of ranks had more inconsistent performance, such as minimap2/SNVer, which for example performed relatively strongly for *N. gonorrhoeae* but poorly for *S. dysenteriae* (Table 2).

**Table 2.** Overall performance of each pipeline per species, calculated as the sum of seven ranks, when reads are aligned to a divergent genome. The seven performance measures for each pipeline (the absolute numbers of true positive, false positive and false negative calls, and the proportion-based precision, recall, F-score, and total error rate per million sequenced bases) are detailed in Supplementary Table 6, with associated ranks in Supplementary Table 7.

While, in general, the accuracy of SNP calling declined with increasing genetic distances, some pipelines were more stable than others (Figure 3B). If considering the median difference in F-score between SNP calls made using the same versus a representative genome, Snippy had smaller differences as the distance between genomes increased (Figure 4).

**Figure 4.**
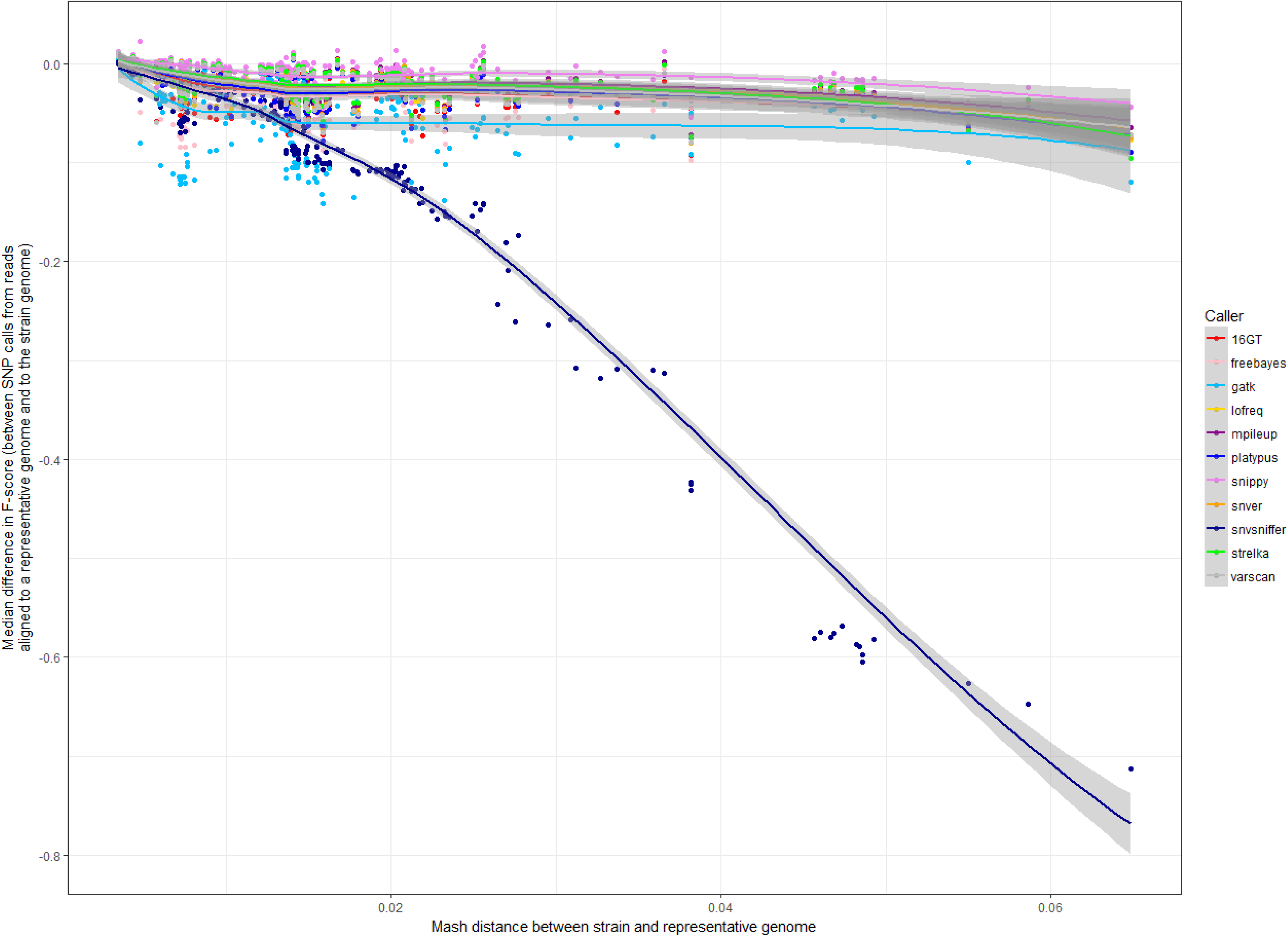
Stability of pipeline performance, in terms of F-score, with increasing genetic distance between the reads and the reference genome. The performance of a SNP calling pipeline decreases with increasing distance between the genome from which reads are sequenced and the reference genome to which they are aligned. Each point shows the median difference in F-score for a pipeline that calls SNPs when the reference genome is the same as the source of the reads, and when it is instead a representative genome for that species. Points are coloured according to the variant caller in each pipeline, with those towards the top of the figure less affected by distance. Lines fitted using LOESS smoothing.

The highest ranked pipelines in Table 2 had small, but practically unimportant, differences in median F-score and so are arguably equivalently strong candidates for a ‘general purpose’ SNP calling solution. For instance, on the basis of F-score alone the performance of Novoalign/mpileup is negligibly different from BWA-mem/mpileup (Figure 5). However, when directly comparing pipelines, similarity of F-score distributions (see Figure 2B) can conceal larger differences in either precision or recall, categorised using the effect size estimator Cliff’s delta [44, 45]. Thus, certain pipelines may be preferred if the aim is to minimise false positive (e.g. for transmission analysis) or maximise true positive (e.g. to identify antimicrobial resistance loci) calls. For instance, although Snippy (the top ranked pipeline in Table 2) is negligibly different from Novoalign/mpileup (the third ranked pipeline) in terms of F-score and precision, the former is more sensitive (Figure 5).

**Figure 5.**
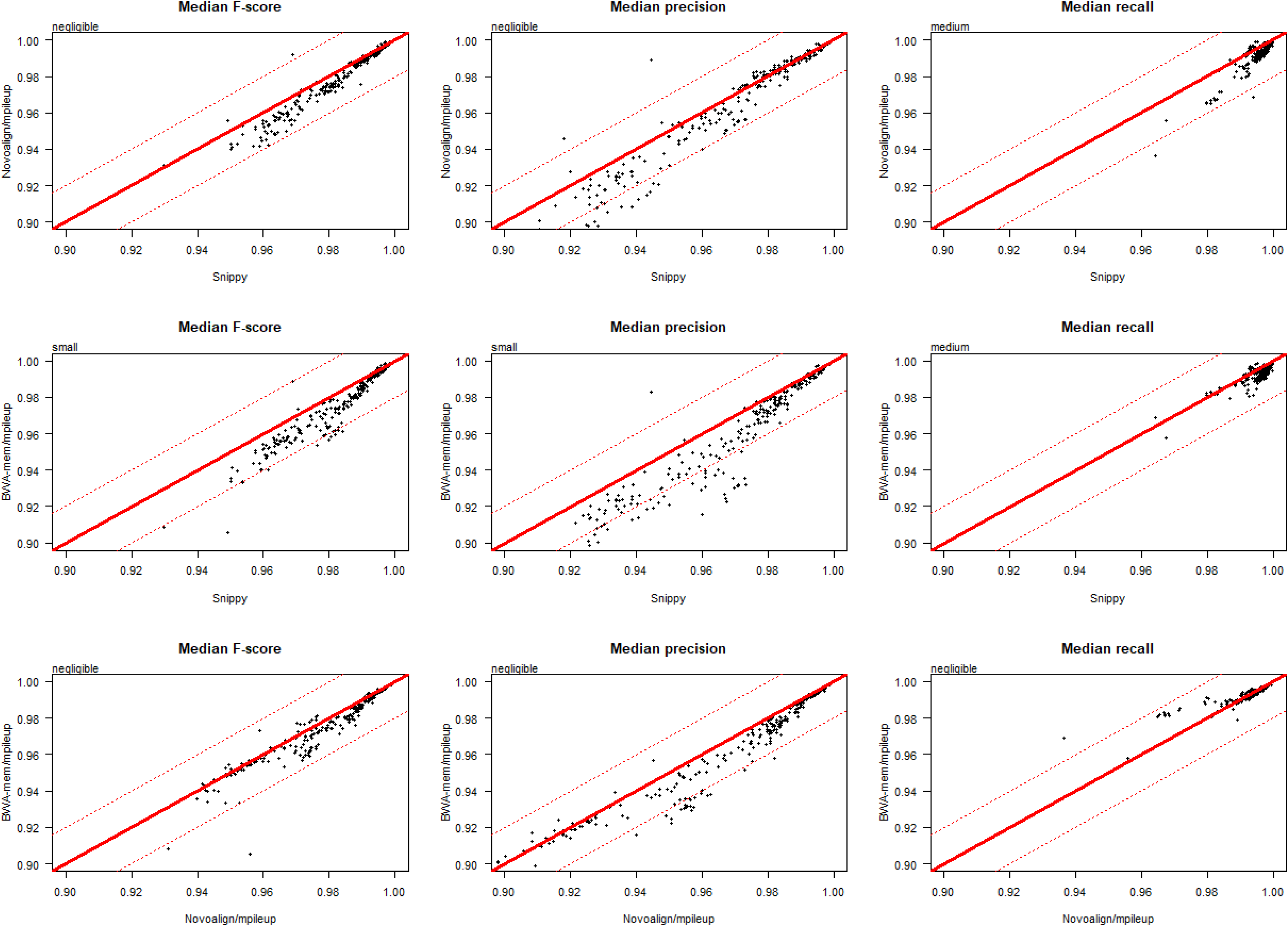
Head-to-head performance comparison of three pipelines, on the basis of precision, recall and F-score. This figure directly compares the performance of three pipelines using simulated data: Snippy, Novoalign/mpileup and BWA/mpileup. Each point indicates the median F-score, precision or recall (columns 1 through 3, respectively), for the genome of one strain per species (n = 254 strains). Raw data for this figure is given in Supplementary Table 6. Text in the top left of each figure is an interpretation of the difference between each pair of distributions, obtained using the R package ‘effsize’ which applies the non-parametric effect size estimator Cliff’s delta to the results of a Mann Whitney U test. An expanded version of this figure, comparing 40 pipelines relative to Snippy, is given as Supplementary Figure 4.

### Comparable accuracy of SNP calling pipelines if using real rather than simulated sequencing data

We used real sequencing data from a previous study comprising 16 environmentally-sourced Gram-negative isolates (all *Enterobacteriaceae*), derived from livestock farms, sewage, and rivers, and cultures of two reference strains (*K. pneumoniae* subsp. *pneumoniae* MGH 78578 and *E. coli* CFT073), for which closed hybrid *de novo* assemblies were generated using both Illumina paired-end short reads and Nanopore long reads [46]. Source locations for each sample, species predictions and NCBI accession numbers are detailed in Supplementary Table 8. The performance statistics for each pipeline are shown in Supplementary Table 9, with an associated ranked summary in Supplementary Table 10.

Lower performance was anticipated for all pipelines, particularly for *Citrobacter* and *Enterobacter* isolates, which had comparatively high Mash distances (> 0.08) between the reads and the representative genome (Supplementary Table 8), far greater than those in the simulations (241 of the 254 simulated genomes had a Mash distance to the representative genome of < 0.04; Supplementary Table 2). Consistent with the simulations (Figure 3A), there was a strong negative correlation between Mash distance and the median F-score across all pipelines (Spearman’s *rho* = −0.83, p = 3.36×10^−5^; Figure 6A), after excluding one prominent outlier (*E. coli* isolate RHB11-C04; see Supplementary Table 8).

**Figure 6.**
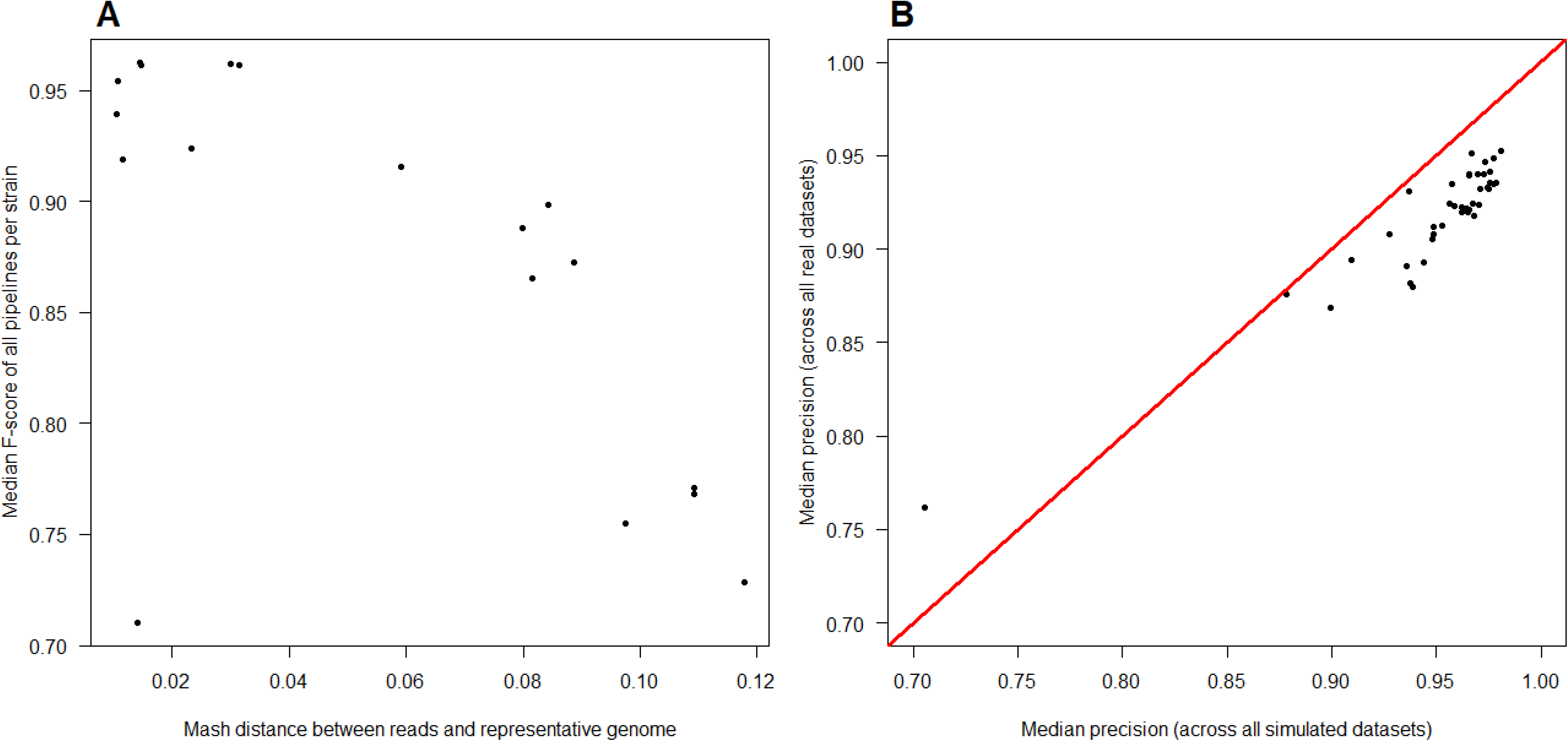
Similarity of performance for pipelines evaluated using both simulated and real sequencing data. Panel A shows that pipelines evaluated using real sequencing data show reduced performance with increasing Mash distances between the reads and the reference genome, similar to that observed with simulated data (see Figure 3A). Each point indicates the median F-score, across all pipelines, for the genome of an environmentally-sourced/reference isolate (detailed in Supplementary Table 8). Panel B shows that pipelines evaluated using real and simulated sequencing data have comparable accuracy. Each point shows the median precision of each of 41 pipelines, calculated across both a divergent set of 254 simulated genomes (2-36 strains from ten clinically common species) and 18 real genomes (isolates of *Citrobacter, Enterobacter, Escherichia* and *Klebsiella*). The outlier pipeline, with lowest precision on both real and simulated data, is Stampy/Freebayes. Raw data for this figure are available in Supplementary Tables 6 (simulated genomes) and 9 (real genomes).

Notably, the median precision of each pipeline, if calculated across the divergent set of simulated genomes, strongly correlated with the median precision calculated across the set of real genomes (Spearman’s *rho* = 0.83, p = 2.81×10^−11^; Figure 6B). While a weaker correlation was seen between simulated and real datasets on the basis of recall (Spearman’s *rho* = 0.41, p = 0.007), this is consistent with the high diversity of *Enterobacteriaceae*, and the accordingly greater number of false negative calls with increased divergence (Supplementary Figure 2).

Overall, this suggests that the accuracy of a given pipeline on simulated data is a reasonable proxy for its performance on real data. While the poorer performing pipelines when using simulated data are similarly poorer performing when using real data, the top ranked pipelines differ, predominantly featuring BWA-mem, rather than Novoalign, as an aligner (Supplementary Table 10). In both cases, however, among the consistently highest performing pipelines is Snippy.

## Discussion

### Reference genome selection strongly affects SNP calling performance

Here we have evaluated 41 SNP calling pipelines, the combination of 4 aligners with 10 callers, plus one self-contained pipeline, Snippy, using reads simulated from 10 clinically relevant species. These reads were first aligned back to their source genome and SNPs called. As expected under these conditions, the majority of SNP calling pipelines showed high precision and sensitivity, although between-species variation was prominent.

We next introduced a degree of divergence between the reference genome and the reads, analogous to having an accurate species-level classification of the reads but no specific knowledge of the strain. For the purposes of this study, we assumed that reference genome selection was essentially arbitrary, equivalent to a community standard representative genome. Such a genome can differ significantly from the sequenced strain, which complicates SNP calling by introducing inter-specific variation between the sequenced reads and the reference. Importantly, all pipelines in this study are expected to perform well if evaluated with human data, i.e. when there is a negligible Mash distance between the reads and the reference. For example, the mean Mash distance between human assembly GRCh38.p12 and the 3 Ashkenazi assemblies of the Genome In A Bottle dataset (deep sequencing of a mother, father and son trio [47–49], available under ENA study accession PRJNA200694 and GenBank assembly accessions GCA_001549595.1, GCA_001549605.1, and GCA_001542345.1, respectively) is 0.001 (i.e., consistent with previous findings that the majority of the human genome has approximately 0.1% sequence divergence [50]). Notably, the highest performing pipeline when reads were aligned to the same genome from which they were simulated, Novoalign/GATK, was also that used by the Genome In A Bottle consortium to align human reads to the reference [47].

While tools initially benchmarked on human data, such as SNVSniffer [34], can in principle also be used on bacterial data, this study shows that in practice many perform poorly. For example, the representative *C. difficile* strain, 630, has a mosaic genome, approximately 11% of which comprises mobile genetic elements [38]. With the exception of reads simulated from *C. difficile* genomes which are erythromycin-sensitive derivatives of 630 (strains 630Derm and 630deltaerm; see [51]), aligning reads to 630 compromises accurate SNP calling, resulting in a lower median F-score across all pipelines (Figure 3A). We also observed similar decreases in F-score for more recombinogenic species such as *N. gonorrhoeae*, which has a phase-variable gene repertoire [52] and has been used to illustrate the ‘fuzzy species’ concept, that recombinogenic bacteria do not form clear and distinct isolate clusters as assayed by phylogenies of common housekeeping loci [53, 54]. By contrast, for clonal species, such as those within the *M. tuberculosis* complex [55], the choice of reference genome has negligible influence on the phylogenetic relationships inferred from SNP calls [56] and, indeed, minimal effect on F-score.

In general, more diverse species have a broader range of Mash distances on Figure 2A (particularly notable for *E. coli*), as do those forming distinct phylogroups, such as the two clusters of *L. monocytogenes*, consistent with the division of this species into multiple primary genetic lineages [57–59].

Therefore, one major finding of this study is that, irrespective of the core components within a SNP calling pipeline, the selection of reference genome has a critical effect on output, particularly for more recombinogenic species. This can to some extent be mitigated by using variant callers that are more robust to increased distances between the reads and the reference, such as Freebayes (employed by Snippy).

A sub-optimal choice of reference genome has previously been shown to result in mapping errors, leading to biases in allelic proportions [60]. Heterologous reference genomes are in general sub-optimal for read mapping, even when there is strict correspondence between orthologous regions, with short reads particularly vulnerable to false positive alignments [61]. There is also an inverse relationship between true positive SNP calls and genetic distance, with a greater number of false positives when the reads diverge from the reference genome [22].

### Study limitations

The experimental design made several simplifying assumptions regarding pipeline usage. Most notably, when evaluating SNP calling when the reference genome diverges from the source of the reads, we needed to convert the coordinates of one genome to those of another, doing so by whole genome alignment. We took a similar approach to that used to evaluate Pilon, an all-in-one tool for correcting draft assemblies and variant calling [62], which made whole genome alignments of the *M. tuberculosis* F11 and H37Rv genomes and used the resulting set of inter-strain variants as a truth set for benchmarking (a method we also used when evaluating each pipeline on real data). While this approach assumes a high degree of contiguity for the whole genome alignment, there are nevertheless significant breaks in synteny between F11 and H37Rv, with two regions deemed particularly hypervariable, in which no variant could be confidently called [62]. For the strain-to-representative genome alignments in this study, we considered SNP calls only within one-to-one alignment blocks and cannot exclude the possibility that repetitive or highly mutable regions within these blocks have been misaligned. However, we did not seek to identify and exclude SNPs from these regions as, even if present, this would have a systematic negative effect on the performance of each pipeline.

Furthermore, when aligning reads from one genome to a different genome, it is not possible to recover all possible SNPs introduced with respect to the former, as some will be found only within genes unique to the original genome (of which there can be many, as bacterial species have considerable genomic diversity; see Supplementary Table 5). Nevertheless, there is a strong relationship between the total number of SNPs introduced *in silico* into one genome and the maximum number of SNPs it is possible to call should reads instead be aligned to a divergent genome (Supplementary Figure 3). In any case, this does not affect the evaluation metrics used for pipeline evaluation, such as F-score, as these are based on proportional relationships of true positive, false positive and false negative calls at variant sites. However, we did not count true negative calls (and thereby assess pipeline specificity) as these can only be made at reference sites, a far greater number of which do not exist when aligning between divergent genomes.

While the programs chosen for this study are in common use and the findings generalisable, it is also important to note that they are a subset of the tools available (see Supplementary Text 1). It is also increasingly common to construct more complex pipelines that call SNPs with one tool and structural variants with another (for example, in [63]). Here, our evaluation concerned only accurate SNP calling, irrespective of the presence of structural variants introduced by sub-optimal reference genome selection (that is, by aligning the reads to a divergent genome) and so does not test dedicated indel calling algorithms. Previous indel-specific variant calling evaluations, using human data, have recommended Platypus [8] or, for calling large indels at low read depths, Pindel [64].

Many of the findings in this evaluation are also based on simulated error-free data for which there was no clear need for pre-processing quality control. While adaptor removal and quality-trimming reads are recommended precautionary steps prior to analysing non-simulated data, previous studies differ as to whether pre-processing increases the accuracy of SNP calls [65], has minimal effect upon them [66], or whether benefits instead depend upon the aligner and reference genome used [22]. While more realistic datasets would be subject to sequencing error, we also expect this to be minimal: Illumina platforms have a per-base error rate < 0.01% [67]. Accordingly, when comparing pipelines taking either error-free or error-containing reads as input, sequencing error had negligible effect on performance (see Supplementary Text 1).

We have also assumed that given the small genome sizes of bacteria, a consistently high depth of coverage is expected in non-simulated datasets, and so have not evaluated pipeline performance on this basis. In any case, a previous study found that with simulated NextSeq reads, variant calling sensitivity was largely unaffected by increases in coverage [40].

### Recommendations for bacterial SNP calling

Our results emphasise that one of the principal difficulties of alignment-based bacterial SNP calling is not pipeline selection *per se* but optimal reference genome selection (or, alternatively, its *de novo* creation, not discussed further). If assuming all input reads are from a single, unknown, origin, then in principle a reference genome could be predicted using a metagenomic classifier such as Centrifuge [68], Kaiju [69] or Kraken [70]. However, correctly identifying the source genome from even a set of single-origin reads is not necessarily simple with the performance of read classifiers depending in large part on the sequence database they query (such as, for instance, EMBL proGenomes [71] or NCBI RefSeq [72]), which can vary widely in scope, redundancy, and degree of curation (see performance evaluations [73, 74]). This is particularly evident among the *Citrobacter* samples in the real dataset, with 3 methods each making different predictions (Supplementary Table 8). Specialist classification tools such as Mykrobe [75] use customised, tightly curated, allele databases and perform highly for certain species (in this case, *M. tuberculosis* and *S. aureus*) although by definition do not have wider utility. An additional complication would also arise from taxonomic disputes such as, for example, *Shigella* spp. being essentially indistinct from *E. coli* [76].

One recommendation, which is quick and simple to apply, would be to test which of a set of candidate reference genomes is most suitable by estimating the distance between each genome and the reads. This can be accomplished using Mash [43], which creates ‘sketches’ of sequence sets (compressed representations of their k-mer distributions) and then estimates the Jaccard index (that is, the fraction of shared k-mers) between each pair of sequences. Mash distances are a proxy both for average nucleotide identity [43] and measures of genetic distance derived from the whole genome alignment of genome pairs (Supplementary Table 2), correlating strongly with the total number of SNPs between the strain genome and the representative genome (Spearman’s *rho* = 0.97, p < 10^−15^), and to a reasonable degree with the proportion of bases unique to the strain genome (Spearman’s *rho* = 0.48, p < 10^−15^). More closely related genomes would have lower Mash distances and so be more suitable as reference genomes for SNP calling. Using a highly divergent genome (such as the representative *Enterobacter* genomes in the real dataset, each of which differs from the reads by a Mash distance > 0.1; Supplementary Table 8) is analogous to variant calling in a highly polymorphic region, such as the human leukocyte antigen, which shows > 10% sequence divergence between haplotypes [50] (i.e., even for pipelines optimised for human data – the majority in this study – this would represent an anomalous use case).

Prior to using Mash (or other sketch-based distance-estimators, such as Dashing [77] or FastANI [78]), broad-spectrum classification tools such as Kraken could be used to narrow down the scope of the search space to a set of fully-sequenced candidate genomes, i.e. those genomes of the taxonomic rank to which the highest proportion of reads could be assigned with confidence.

In the future, reads from long-read sequencing platforms, such as Oxford Nanopore, are less likely to be ambiguously mapped within a genomic database and so in principle are simpler to classify (sequencing error rate notwithstanding), making it easier to select a suitable reference genome. However, long-read platforms can also, in principle if not yet routinely, generate complete *de novo* bacterial genomes [79] for downstream SNP calling, possibly removing the need to choose a reference entirely. Similarly, using a reference pan-genome instead of a singular representative genome could also maximise the number of SNP calls by reducing the number of genes not present in the reference [80].

If considering the overall performance of a pipeline as the sum of the 7 different ranks for the different metrics considered, then averaged across the full set of species’ genomes, the highest performing pipelines are, with simulated data, Snippy and those utilising Novoalign in conjunction with LoFreq or mpileup (Table 2), and with real data, Snippy and those utilising BWA-mem in conjunction with Strelka or mpileup (Supplementary Table 10).

Some of the higher-performing tools apply error-correction models that also appear suited to bacterial datasets with high SNP density, despite their original primary use case being in different circumstances. For instance, SNVer (which in conjunction with BWA-mem, ranks second to Snippy for *N. gonorrhoeae;* see Table 2) implements a statistical model for calling SNPs from pooled DNA samples, where variant allele frequencies are not expected to be either 0, 0.5 or 1 [33]. SNP calling from heterogeneous bacterial populations with high mutation rates, in which only a proportion of cells may contain a given mutation, is also conceptually similar to somatic variant calling in human tumours, where considerable noise is expected [60] (this is a recommended use case for Strelka, which performed highly on real data; Supplementary Table 10).

Irrespective of pipeline employed, increasing Mash distances between the reads and the reference increases the number of false negative calls (Supplementary Figure 2). Nevertheless, Snippy, which employs Freebayes, is particularly robust to this, being among the most sensitive pipelines (Figure 5 and Supplementary Figure 4). Notably, Freebayes is haplotype-based, calling variants based on the literal sequence of reads aligned to a particular location, so avoiding the problem of one read having multiple possible alignments (increasingly likely with increasing genomic diversity) but only being assigned to one of them. However, as distance increases further, it is likely that reads will cease being misaligned (which would otherwise increase the number of false positive calls) but rather they will not be aligned at all, being too dissimilar to the reference genome.

With an appropriate selection of reference genome, many of these higher-performing pipelines could be optimised to converge on similar results by tuning parameters and post-processing VCFs with specific filtering criteria, another routine task for which there are many different choices of application [81–84]. In this respect, the results of this study should be interpreted as a range-finding exercise, drawing attention to those SNP calling pipelines which, under default conditions, are generally higher-performing and which may be most straightforwardly optimised to meet user requirements.

## Conclusions

We have performed a comparison of SNP calling pipelines across both simulated and real data in multiple bacterial species, allowing us to benchmark their performance for this specific use. We find that all pipelines show extensive species-specific variation in performance, which has not been apparent from the majority of existing, human-centred, benchmarking studies. While aligning to a single representative genome is common practice in eukaryotic SNP calling, in bacteria the sequence of this genome may diverge considerably from the sequence of the reads. A critical factor affecting the accuracy of SNP calling is thus the selection of a reference genome for alignment. This is complicated by ambiguity as to the strain of origin for a given set of reads, which is perhaps inevitable for many recombinogenic species, a consequence of the absence (or impossibility) of a universal species concept for bacteria. For many clinically common species, excepting *M. tuberculosis*, the use of standard ‘representative’ reference genomes can compromise accurate SNP calling by disregarding genomic diversity. By first considering the Mash distance between the reads and a candidate set of reference genomes, a genome with minimal distance may be chosen that, in conjunction with one of the higher performing pipelines, can maximise the number of true variants called.

## Materials and Methods

### Simulating truth sets of SNPs for pipeline evaluation

264 genomes, representing a range of strains from 10 bacterial species, and their associated annotations, were obtained from the NCBI Genome database [85] (https://www.ncbi.nlm.nih.gov/genome, accessed 16^th^ August 2018), as detailed in Supplementary Table 2. One genome per species is considered to be a representative genome (criteria detailed at https://www.ncbi.nlm.nih.gov/refseq/about/prokaryotes/, accessed 16^th^ August 2018), indicated in Supplementary Table 2. Strains with incomplete genomes (that is, assembled only to the contig or scaffold level) or incomplete annotations (that is, with no associated GFF, necessary to obtain gene coordinates) were excluded, as were those with multiple available genomes (that is, the strain name was not unique). After applying these filters, all species were represented by approx. 30 complete genomes (28 *C. difficile*, 29 *M. tuberculosis* and 36 *S. pneumoniae*), with the exceptions of *N. gonorrhoeae* (n = 15) and *S. dysenteriae* (n = 2). For the 5 remaining species (*E. coli, K. pneumoniae, L. monocytogenes*, *S. aureus* and *S. enterica*), there are > 100 usable genomes each. As it was not computationally tractable to test every genome, we chose a subset of isolates based on stratified selection by population structure. We created all-against-all distance matrices using the ‘triangle’ component of Mash v2.1 [43], then constructed dendrograms (Supplementary Figures 5 to 9) from each matrix using the neighbour joining method, as implemented in MEGA v7.0.14 [86]. By manually reviewing the topology, 30 isolates were chosen per species to create a representative sample of its diversity.

For each genome used in this study, we excluded, if present, any non-chromosomal (i.e. circular plasmid) sequence. A simulated version of each core genome, with exactly 5 randomly generated SNPs per genic region, was created using Simulome v1.2 [87] with parameters --whole_genome=TRUE --snp=TRUE --num_snp=5. As the coordinates of some genes overlap, not all genes will contain simulated SNPs. The number of SNPs introduced into each genome (from approximately 8000 to 25,000) and the median distance between SNPs (from approximately 60 to 120 bases) is detailed in Supplementary Table 2.

The coordinates of each SNP inserted into a given genome are, by definition, genome- (that is, strain-) specific. As such, it is straightforward to evaluate pipeline performance when reads from one genome are aligned to the same reference. However, in order to evaluate pipeline performance when reads from one genome are aligned to the genome of a divergent strain (that is, the representative genome of that species), the coordinates of each strain’s genome need to be converted to representative genome coordinates. To do so, we made whole genome (core) alignments of the representative genome to both versions of the strain genome (one with and one without SNPs introduced *in silico*) using nucmer and dnadiff, components of MUMmer v4.0.0beta2 [41], with default parameters (illustrated in Figure 1). For one-to-one alignment blocks, differences between each pair of genomes were identified using MUMmer show-snps with parameters −Clr −x 1, with the tabular output of this program converted to VCF by the script MUMmerSNPs2VCF.py (https://github.com/liangjiaoxue/PythonNGSTools, accessed 16^th^ August 2018). The two resulting VCFs contain the location of all SNPs relative to the representative genome (i.e. inclusive of those introduced *in silico*), and all inter-strain variants, respectively. We excluded from further analysis two strains with poor-quality strain-to-representative whole genome alignments, both calling < 10% of the strain-specific *in silico* SNPs (Supplementary Table 11). The proportion of *in silico* SNPs recovered by whole genome alignment is detailed in Supplementary Table 11 and is, in general, high: of the 254 whole genome alignments of non-representative to representative strains across the 10 species, 222 detect > 80% of the *in silico* SNPs and 83 detect > 90%. For the purposes of evaluating SNP calling pipelines when the reference genome differs from the reads, we are concerned only with calling the truth set of *in silico* SNPs and so discard inter-strain variants (see below). More formally, when using each pipeline to align reads to a divergent genome, we are assessing the concordance of its set of SNP calls with the set of nucmer calls. However, it is possible that for a given call, one or more of the pipelines are correct and nucmer is incorrect. To reduce this possibility, a parallel set of whole genome alignments were made using Parsnp v1.2 with default parameters [42], with the exported SNPs contrasted with the nucmer VCF.

Thus, when aligning to a divergent genome, the truth set of *in silico* SNPs (for which each pipeline is scored for true positives) are those calls independently identified by both nucmer and Parsnp. Similarly, the set of inter-strain positions are those calls made by one or both of nucmer and Parsnp. As we are not concerned with the correctness of these calls, the lack of agreement between the two tools is not considered further; rather, this establishes a set of ambiguous positions which are discarded when VCFs are parsed.

Simulated SNP-containing genomes, sets of strain-to-representative genome SNP calls (made by both nucmer and Parsnp), and the final truth sets of SNPs are available in Supplementary Dataset 1 (hosted online via the Oxford Research Archive at http://dx.doi.org/10.5287/bodleian:AmNXrjYN8).

### Evaluating SNP calling pipelines using simulated data

From each of 254 SNP-containing genomes, 3 sets of 150bp and 3 sets of 300bp paired-end were simulated using wgsim, a component of SAMtools v1.7 [21]. This requires an estimate of average insert size (the length of DNA between the adapter sequences), which in real data is often variable, being sensitive to the concentration of DNA used [88]. For read length *x*, we assumed an insert size of 2.2x, i.e. for 300bp reads, the insert size is 660bp (Illumina paired-end reads typically have an insert longer than the combined length of both reads [89]). The number of reads simulated from each genome is detailed in Supplementary Table 3 and is equivalent to a mean 50-fold base-level coverage, i.e. (50 x genome length)/read length.

Perfect (error-free) reads were simulated from each SNP-containing genome using wgsim parameters -e 0 -r 0 -R 0 -X 0 -A 0 (respectively, the sequencing error rate, mutation rate, fraction of indels, probability an indel is extended, and the fraction of ambiguous bases allowed).

Each set of reads was then aligned both to the genome of the same strain and to the representative genome of that species (from which the strain will diverge), with SNPs called using 41 different SNP calling pipelines (10 callers each paired with 4 aligners, plus the self-contained Snippy). The programs used, including version numbers and sources, are detailed in Supplementary Table 1, with associated command lines in Supplementary Text 1. All pipelines were run using a high-performance cluster employing the Open Grid Scheduler batch system on Scientific Linux 7. No formal assessment was made of pipeline run time or memory usage. This was because given the number of simulations it was not tractable to benchmark run time using, for instance, a single core. The majority of programs in this study permit multithreading (all except the callers 16GT, GATK, Platypus, SNVer, and SNVSniffer) and so are in principle capable of running very rapidly. We did not seek to optimise each tool for any given species and so made only a minimum effort application of each pipeline, using default parameters and minimal VCF filtering (see below). This is so that we obtain the maximum possible number of true positives from each pipeline under reasonable use conditions.

While each pipeline comprises one aligner and one caller, there are several ancillary steps common in all cases. After aligning reads to each reference genome, all BAM files were cleaned, sorted, had duplicate reads marked and were indexed using Picard Tools v2.17.11 [90] CleanSam, SortSam, MarkDuplicates and BuildBamIndex, respectively. We did not add a post-processing step of local indel realignment (common in older evaluations, e.g., [12]) as this had negligible effect upon pipeline performance, with many variant callers (including GATK HaplotypeCaller and Freebayes) already incorporating a method of haplotype assembly (see Supplementary Text 1).

Each pipeline produces a VCF as its final output. As with a previous evaluation [26], all VCFs were regularised using the vcfallelicprimitives module of vcflib v1.0.0-rc2 (https://github.com/ekg/vcflib), so that different representations of the same indel or complex variant were not counted separately (these variants can otherwise be presented correctly in multiple ways). This module splits adjacent SNPs into individual SNPs, left-aligns indels and regularizes the representation of complex variants.

Different variant callers populate their output VCFs with different contextual information. Before evaluating the performance of each pipeline, all regularised VCFs were subject to minimal parsing to retain only high-confidence variants. This is because many tools record variant sites even if they have a low probability of variation, under the reasonable expectation of parsing. Some pipelines (notably Snippy) apply their own internal set of VCF filtering criteria, giving the user the option of a ‘raw’ or ‘filtered’ VCF; in such cases, we retain the filtered VCF as the default recommendation. Where possible, (additional) filter criteria were applied as previously used by, and empirically selected for, COMPASS (Complete Pathogen Sequencing Solution; https://github.com/oxfordmmm/CompassCompact), an analytic pipeline employing Stampy and mpileup for base calling non-repetitive core genome sites (outlined in Supplementary Text 1 with filter criteria described in [91] and broadly similar to those recommended by a previous study for maximising SNP validation rate [92]). No set of generic VCF hard filters can be uniformly applied because each caller quantifies different metrics (such as the number of forward and reverse reads supporting a given call) and/or reports the outcome of a different set of statistical tests, making filtering suggestions on this basis. For instance, in particular circumstances, GATK suggests filtering on the basis of the fields ‘FS’, ‘MQRankSum’ and ‘ReadPosRankSum’, which are unique to it (detailed at https://software.broadinstitute.org/gatk/documentation/article.php?id=6925, accessed 2^nd^ April 2019). Where the relevant information was included in the VCF, SNPs were required to have (a) a minimum Phred score of 20, (b) > 5 reads mapped at that position, (c) at least one read in each direction in support of the variant, and (d) >75% of reads supporting the alternative allele. These criteria were implemented with the ‘filter’ module of BCFtools v1.7 [21] using parameters detailed in Supplementary Table 12.

From these filtered VCFs, evaluation metrics were calculated as detailed below.

### Evaluating SNP calling pipelines using real sequencing data

Parallel sets of 150 bp Illumina HiSeq 4000 paired-end short reads and ONT long reads were obtained from 16 environmentally-sourced samples from the REHAB project (‘the environmental REsistome: confluence of Human and Animal Biota in antibiotic resistance spread’; http://modmedmicro.nsms.ox.ac.uk/rehab/), as detailed in [46]: 4 *Enterobacter* spp., 4 *Klebsiella* spp., 4 *Citrobacter* spp., and 4 *Escherichia coli*, with species identified using MALDI-TOF (matrix-assisted laser desorption ionization time-of-flight) mass spectrometry, plus sub-cultures of stocks of two reference strains *K. pneumoniae* subsp. *pneumoniae* MGH 78578 and *E. coli* CFT073. Additional predictions were made using both the protein- and nucleotide-level classification tools Kaiju v1.6.1 [69] and Kraken2 v2.0.7 [93], respectively. Kaiju was used with two databases, one broad and one deep, both created on 5^th^ February 2019: ‘P’ (http://kaiju.binf.ku.dk/database/kaiju_db_progenomes_2019-02-05.tgz; > 20 million bacterial and archaeal genomes from the compact, manually curated, EMBL proGenomes [94], supplemented by approximately 10,000 viral genomes from NCBI RefSeq [95]) and ‘E’ (http://kaiju.binf.ku.dk/database/kaiju_db_nr_euk_2019-02-05.tgz; > 100 million bacterial, archaeal, viral and fungal genomes from NCBI nr, alongside various microbial eukaryotic taxa). Kaiju was run with parameters -e 5 and -E 0.05 which, respectively, allow 5 mismatches per read and filter results on the basis of an E-value threshold of 0.05. The read classifications from both databases were integrated using the Kaiju ‘mergeOutputs’ module, which adjudicates based on the lowest taxonomic rank of each pair of classifications, provided they are within the same lineage, else re-classifies the read at the lowest common taxonomic rank ancestral to the two. Kraken2 was run with default parameters using the MiniKraken2 v1 database (https://ccb.jhu.edu/software/kraken2/dl/minikraken2_v1_8GB.tgz, created 12^th^ October 2018), which was built from the complete set of NCBI RefSeq bacterial, archaeal and viral genomes.

Hybrid assemblies were produced using methods detailed in [46] and briefly recapitulated here. Illumina reads were processed using COMPASS (see above). ONT reads were adapter-trimmed using Porechop v0.2.2 (https://github.com/rrwick/Porechop) with default parameters, and then error-corrected and sub-sampled (preferentially selecting the longest reads) to 30-40x coverage using Canu v1.5 [96] with default parameters. Finally, Illumina-ONT hybrid assemblies for each genome were generated using Unicycler v0.4.0 [39] with default parameters. The original study found high agreement between these assemblies and those produced using hybrid assembly with PacBio long reads rather than ONT, giving us high confidence in their robustness.

In the simulated datasets, SNPs are introduced *in silico* into a genome, with reads containing these SNPs then simulated from it. With this dataset, however, there are no SNPs within each genome: we have only the short reads (that is, real output from an Illumina sequencer) and the genome assembled from them (with which there is an expectation of near-perfect read mapping).

To evaluate pipeline performance when the reads are aligned to a divergent genome, reference genomes were selected as representative of the predicted species, with distances between the two calculated using Mash v2.1 [43] and spanning approximately equal intervals from 0.01 to 0.12 (representative genomes and Mash distances are detailed in Supplementary Table 8). The truth set of SNPs between the representative genome and each hybrid assembly was the intersection of nucmer and Parsnp calls, as above.

Samples, source locations, MALDI ID scores and associated species predictions are detailed in Supplementary Table 8. Raw sequencing data and assemblies have been deposited with the NCBI under BioProject accession PRJNA42251 (https://www.ncbi.nlm.nih.gov/bioproject/PRJNA422511).

### Evaluation metrics

For each pipeline, we calculated the absolute number of true positive (TP; the variant is in the simulated genome and correctly called by the pipeline), false positive (FP; the pipeline calls a variant which is not in the simulated genome) and false negative SNP calls (FN; the variant is in the simulated genome but the pipeline does not call it). We did not calculate true negative calls for two reasons. Firstly, to do so requires a VCF containing calls for all sites, a function offered by some variant callers (such as mpileup) but not all. Secondly, when aligning reads to a divergent genome, a disproportionately large number of reference sites will be excluded, particularly in more diverse species (for example, gene numbers in *N. gonorrhoeae* differ by up to a third; see Supplementary Table 5).

We then calculated the precision (positive predictive value) of each pipeline as TP/(TP+FP), recall (sensitivity) as TP/(TP+FN), miss rate as FN/(TP+FN), and total number of errors (FP+FN) per million sequenced bases. We did not calculate specificity as this depends on true negative calls. We also calculated the F-score (as in [40]), which considers precision and recall with equal weight: F = 2 * ((precision * recall) / (precision + recall)). The F-score evaluates each pipeline as a single value bounded between 0 and 1 (perfect precision and recall). We also ranked each pipeline based on each metric so that – for example – the pipeline with the highest F-score, and the pipeline with the lowest number of false positives, would be rank 1 in their respective distributions. As an additional ‘overall performance’ measure, we calculated the sum of ranks for the 7 core evaluation metrics (the absolute numbers of TP, FP and FN calls, and the proportion-based precision, recall, F-score, and total error rate per million sequenced bases). Pipelines with a lower sum of ranks would, in general, have higher overall performance.

We note that when SNPs are called after aligning reads from one strain to that of a divergent strain, the SNP calling pipeline will call positions for both the truth set of strain-specific *in silico* SNPs and any inter-strain variants. To allow a comparable evaluation of pipelines in this circumstance, inter-strain calls (obtained using nucmer and Parsnp; see above) are discarded and not explicitly considered either true positive, false positive or false negative. While the set of true SNPs when aligning to a divergent strain will be smaller than that when aligned to the same strain (because all SNPs are simulated in genic regions but not all genes are shared between strains), this will not affect proportion-based evaluation metrics, such as F-score.

### Effect size of differences in the F-score distribution between pipelines

Differences between distributions are assessed by Mann Whitney U tests, with results interpreted using the non-parametric effect size estimator Cliff’s delta [44, 45], estimated at a confidence level of 95% using the R package effsize v0.7.1 [97]. Cliff’s delta employs the concept of dominance (which refers to the degree of overlap between distributions) and so is more robust when distributions are skewed. Estimates of delta are bound in the interval (−1, 1), with extreme values indicating a lack of overlap between groups (respectively, set 1 ≪ set 2 and set 1 ≫ set 2). Distributions with |delta| < 0.147 are negligibly different, as in [98]. Conversely, distributions with |delta| >= 0.60 are considered to have large differences.

## Supplementary Tables

**Supplementary Table 1.** Sources of software.

**Supplementary Table 2.** Genomes into which SNPs were introduced *in silico*, and various measures of distance between each strain’s genome and the representative genome of that species.

**Supplementary Table 3.** Summary statistics of SNP calling pipelines after aligning reads to the same reference genome as their origin.

**Supplementary Table 4.** Ranked performance of SNP calling pipelines after aligning reads to the same reference genome as their origin.

**Supplementary Table 5.** Genome size diversity within 5 clinically common bacterial species.

**Supplementary Table 6.** Summary statistics of SNP calling pipelines after aligning reads to a reference genome differing from their origin.

**Supplementary Table 7.** Ranked performance of SNP calling pipelines after aligning reads to reference genome differing from their origin.

**Supplementary Table 8.** Environmentally-sourced/reference Gram-negative isolates and associated representative genomes.

**Supplementary Table 9.** Summary statistics of SNP calling pipelines after aligning real reads to a reference genome differing from their origin.

**Supplementary Table 10.** Ranked performance of SNP calling pipelines after aligning real reads to reference genome differing from their origin.

**Supplementary Table 11.** Proportion of strain-specific *in silico* SNPs detected in whole genome alignments between the strain genome and a representative genome.

**Supplementary Table 12.** VCF filtering parameters, as used by BCFtools.

**Supplementary Table 13.** Summary statistics of SNP calling pipelines after aligning both error-free and error-containing reads to the same reference genome as their origin.

**Supplementary Table 14.** Summary statistics of SNP calling pipelines after aligning both error-free and error-containing reads to a reference genome differing from their origin.

**Supplementary Table 15.** Summary statistics of SNP calling pipelines after aligning error-free reads to a reference genome differing from their origin, both with and without local indel realignment.

## Supplementary Figures

**Supplementary Figure 1. Reduced performance of SNP calling pipelines with increasing genetic distance between the reads and the reference genome (assayed as total number of SNPs).**

The median F-score across a set of 41 pipelines, per strain, decreases as the distance between the strain and the reference genome increases (assayed as the total number of SNPs between the strain and representative genome, i.e. the set of strain-specific *in silico* SNPs plus inter-strain SNPs). Each point indicates the genome of one strain per species (n = 254 strains). Points are coloured by the species of each strain (n = 10 species). Summary statistics for each pipeline are shown in Supplementary Table 6, performance ranks in Supplementary Table 7 and the genetic distance between strains in Supplementary Table 2. Quantitatively similar results are seen if assaying distance as the Mash distance, which is based on the proportion of k-mers shared between genomes (Figure 3A).

**Supplementary Figure 2. Decreasing sensitivity (that is, an increased number of false negative calls) with increasing genetic distance between the reads and the reference genome (assayed as Mash distance).**

The median sensitivity (recall) across a set of 41 pipelines, per strain, increases as the distance between the strain and the reference genome increases (assayed as the Mash distance, which is based on the proportion of shared k-mers between genomes). Each point indicates the genome of one strain per species (n = 254 strains). Points are coloured by the species of each strain (n = 10 species). Summary statistics for each pipeline are shown in Supplementary Table 6, performance ranks in Supplementary Table 7 and the genetic distance between strains in Supplementary Table 2.

**Supplementary Figure 3. Total number of SNPs it is possible to call should reads from one strain be aligned to a representative genome of that species.**

Strong correlation between the total number of SNPs introduced *in silico* into one genome and the maximum number of SNPs it is possible to call assuming reads from the former are aligned to a representative genome of that species (which will not necessarily contain the same complement of genes). Each point represents the genome of one strain, with genomes detailed in Supplementary Table 2. The line y = x is shown in red.

**Supplementary Figure 4. Head-to-head performance comparison of all pipelines relative to Snippy, on the basis of F-score.**

This figure directly compares the performance, using simulated data, of 40 pipelines relative to Snippy. Each point indicates the median F-score for the genome of one strain per species (n = 254 strains). Data for Snippy is plotted on the x-axis, and for the named pipeline on the y-axis. Raw data for this figure is given in Supplementary Table 6. Text in the top left of each figure is an interpretation of the difference between each pair of distributions, obtained using the R package ‘effsize’ which applies the non-parametric effect size estimator Cliff’s delta to the results of a Mann Whitney U test.

**Supplementary Figure 5. Selection of *E. coli* isolates by manual review of dendrogram topology.**

There are numerous usable complete genomes for *E. coli*. For the SNP calling evaluation, a subset of isolates was selected (indicated in red boxes) so as to maximise the diversity of clades represented. To do so, an all-against-all distance matrix for each genome was created using the ‘triangle’ component of Mash v2.1, with a dendrogram constructed using the neighbour joining method implemented in MEGA v7.0.14. Sources for the selected genomes are given in Supplementary Table 2.

**Supplementary Figure 6. Selection of *K. pneumoniae* isolates by manual review of dendrogram topology.**

There are numerous usable complete genomes for *K. pneumoniae*. For the SNP calling evaluation, a subset of isolates was selected (indicated in red boxes) so as to maximise the diversity of clades represented. To do so, an all-against-all distance matrix for each genome was created using the ‘triangle’ component of Mash v2.1, with a dendrogram constructed using the neighbour joining method implemented in MEGA v7.0.14. Sources for the selected genomes are given in Supplementary Table 2.

**Supplementary Figure 7. Selection of *L. monocytogenes* isolates by manual review of dendrogram topology.**

There are numerous usable complete genomes for *L. monocytogenes*. For the SNP calling evaluation, a subset of isolates was selected (indicated in red boxes) so as to maximise the diversity of clades represented. To do so, an all-against-all distance matrix for each genome was created using the ‘triangle’ component of Mash v2.1, with a dendrogram constructed using the neighbour joining method implemented in MEGA v7.0.14. Sources for the selected genomes are given in Supplementary Table 2.

**Supplementary Figure 8. Selection of *S. enterica* isolates by manual review of dendrogram topology.**

There are numerous usable complete genomes for *S. enterica*. For the SNP calling evaluation, a subset of isolates was selected (indicated in red boxes) so as to maximise the diversity of clades represented. To do so, an all-against-all distance matrix for each genome was created using the ‘triangle’ component of Mash v2.1, with a dendrogram constructed using the neighbour joining method implemented in MEGA v7.0.14. Sources for the selected genomes are given in Supplementary Table 2.

**Supplementary Figure 9. Selection of *S. aureus* isolates by manual review of dendrogram topology.**

There are numerous usable complete genomes for *S. aureus*. For the SNP calling evaluation, a subset of isolates was selected (indicated in red boxes) so as to maximise the diversity of clades represented. To do so, an all-against-all distance matrix for each genome was created using the ‘triangle’ component of Mash v2.1, with a dendrogram constructed using the neighbour joining method implemented in MEGA v7.0.14. Sources for the selected genomes are given in Supplementary Table 2.

## Supplementary Datasets

**Supplementary Dataset 1. Simulated datasets for evaluating bacterial SNP calling pipelines.**

This archive contains the set of 254 SNP-containing genomes, VCFs containing the nucmer and Parsnp strain-to-representative genome SNP calls, and the final truth sets of SNPs used for evaluation.

## Declarations

## Ethics approval and consent to participate

Not applicable.

## Consent for publication

Not applicable.

## Availability of data and material

All data analysed during this study are included in this published article and its supplementary information files. The simulated datasets generated during this study – comprising the SNP-containing genomes, log files of the SNPs introduced into each genome, and VCFs of strain-to-representative genome SNP calls – are available in Supplementary Dataset 1 (hosted online via the Oxford Research Archive at http://dx.doi.org/10.5287/bodleian:AmNXrjYN8). Raw sequencing data and assemblies from the REHAB project, described in [46], are available in the NCBI under BioProject accession PRJNA42251 (https://www.ncbi.nlm.nih.gov/bioproject/PRJNA422511).

## Authors’ contributions

SJB conceived of and designed the study with support from DF, DWE, TEAP, DWC and ASW. SJB performed all informatic analyses related to the SNP calling evaluation. ELC contributed to the acquisition of data and computational resources. NDM, LPS and NS generated and provided the reads and assemblies comprising the REHAB sequencing dataset. LPS created Figure 1. SJB wrote the manuscript, with edits from all other authors. All authors read and approved the final manuscript.

## Acknowledgements

The authors would also like to thank the REHAB consortium, which currently includes (bracketed individuals in the main author list): Abuoun M, Anjum M, Bailey MJ, Barker L, Brett H, Bowes MJ, Chau K, (Crook DW), (De Maio N), Gilson D, Gweon HS, Hubbard ATM, Hoosdally S, Kavanagh J, Jones H, (Peto TEA), Read DS, Sebra R, (Shaw LP), Sheppard AE, Smith R, (Stoesser N), Stubberfield E, Swann J, (Walker AS), Woodford N.

