## Supplementary Text 1 for "Genomic diversity affects the accuracy of bacterial SNP calling pipelines"

This supplementary text file details (a) the command lines used when evaluating each aligner/caller combination, and elaborates on (b) the selection criteria by which programs were excluded from this study, (c) the rationale for simulating error-free reads, (d) the rationale for excluding a local indel realignment step, and (e) the filter criteria used to parse each VCF.

(1) **Command lines**

Command lines for each aligner/caller combination are detailed below. For the purpose of this evaluation, Perl scripts were used to generate each set of command lines, and accordingly, they are presented in this fashion. The colour-coded text refers to variables which may be interpolated. For instance, **$ref** refers to the reference genome (or its appropriate index) used as input to each aligner/caller, and **$fq_1** and **$fq_2** to the paired-end fastq files, respectively.

Output filenames were also given a standard nomenclature, comprising a **$root** component (specifying the genome, read length [150 or 300 bp] and replicate number), followed by the **$aligner**, **$caller** and file type suffix, for example “**$root.$aligner.$caller**.vcf”. Intermediary files may be identified by text immediately preceding the suffix, such as “**$root.$aligner.$caller**.unsorted.bam”.

The exception to the following set of combinatorial aligner/caller command lines are those used for the self-contained Snippy:

snippy --cpus 40 --outdir **$root** --prefix **$root** --cleanup --ref **$ref** --R1 **$fq_1** --R2 **$fq_2**

mv **$root/$root**.filt.vcf **$root.$aligner.$caller**.vcf

rm -r **$root**

Note that this produces a VCF, **$root/$root**.filt.vcf, to which Snippy has already applied its own internal set of filter criteria, on the basis of, for BAM parsing, minimum read mapping quality (default: Phred score ≥ 60) and minimum base quality (default: Phred score ≥ 13), and, for VCF parsing, minimum site depth (default: ≥ 10 reads) and minimum variant quality (default: Phred score ≥ 100). The latter are more stringent than the minimum site depth and variant quality thresholds applied to filter the final VCF produced by each pipeline (see below).

**First step of each pipeline: command lines for alignment**

All command lines for each of the 4 aligners (BWA, minimap2, Novoalign, and Stampy) produce as output a Picard-cleaned, but unsorted, BAM: **$root.$aligner**.unsorted.bam.

BWA

bwa mem -R '@RG\tID:group\tSM:sample\tPL:Illumina\tLIB:lib\tPU:unit' -t 40 -M **$ref $fq_1 $fq_2** | samtools view -Shb - > **$root.$aligner**.unsorted.bam

java -jar picard.jar CleanSam INPUT=**$root.$aligner**.unsorted.bam OUTPUT=**$root.$aligner**.unsorted.cleaned.bam

minimap2

minimap2 -ax sr **$ref** -R '@RG\tID:group\tSM:sample\tPL:Illumina\tLIB:lib\tPU:unit' -t 40 $fq_1 $fq_2 | samtools view -Shb - > **$root.$aligner**.unsorted.bam

java -jar picard.jar CleanSam INPUT=**$root.$aligner**.unsorted.bam OUTPUT=**$root.$aligner**.unsorted.cleaned.bam

rm **$root.$aligner**.unsorted.bam

mv **$root.$aligner**.unsorted.cleaned.bam **$root.$aligner**.unsorted.bam

Novoalign

novoalign -c 40 -d **$ref** -F STDFQ -o SAM $'@RG\tID:group\tSM:sample\tPL:Illumina\tLIB:lib\tPU:unit' -f **$fq_1 $fq_2** > **$root.$aligner**.unsorted.sam

samtools view -bS **$root.$aligner**.unsorted.sam > **$root.$aligner**.unsorted.bam

rm **$root.$aligner**.unsorted.sam

java -jar picard.jar CleanSam INPUT=**$root.$aligner**.unsorted.bam OUTPUT=**$root.$aligner**.unsorted.cleaned.bam

rm **$root.$aligner**.unsorted.bam

mv **$root.$aligner**.unsorted.cleaned.bam **$root.$aligner**.unsorted.bam

Stampy

stampy.py -g **$ref** -h **$ref** -t 40 -M **$fq_1 $fq_2** | samtools view -Sb - > **$root.$aligner**.unsorted.no_read_groups.bam

java -jar picard.jar CleanSam INPUT=**$root.$aligner**.unsorted.no_read_groups.bam OUTPUT=**$root.$aligner**.unsorted.no_read_groups.cleaned.bam

rm **$root.$aligner**.unsorted.no_read_groups.bam

java -jar picard.jar AddOrReplaceReadGroups INPUT=**$root.$aligner**.unsorted.no_read_groups.cleaned.bam OUTPUT=**$root.$aligner**.unsorted.bam RGID=**$root** RGLB=lib RGPL=Illumina RGPU=unit RGSM=sample

rm **$root.$aligner**.unsorted.no_read_groups.cleaned.bam

**Second step of each pipeline: command lines for post-processing BAMs**

These command lines constitute an intermediary step in each pipeline, taking as input an unsorted BAM, **$root.$aligner**.unsorted.bam, and producing as output a sorted, de-duplicated and indexed BAM, **$root.$aligner**.bam. This is the final output of the alignment process and used as input to the third step of each pipeline: variant calling. All intermediary BAMs are discarded.

java -jar picard.jar SortSam INPUT=**$root.$aligner**.unsorted.bam OUTPUT=**$root.$aligner**.sorted.bam SORT_ORDER=coordinate

java -jar picard.jar MarkDuplicates INPUT=**$root.$aligner**.sorted.bam OUTPUT=**$root.$aligner**.bam METRICS_FILE=**$root.$aligner**.metrics ASSUME_SORTED=true

java -jar picard.jar BuildBamIndex INPUT=**$root.$aligner**.bam

rm **$root.$aligner**.unsorted.bam **$root.$aligner**.sorted.bam **$root.$aligner**.metrics

**Final step of each pipeline: command lines for variant calling**

All command lines for each of the 10 aligners (16GT, Freebayes, GATK HaplotypeCaller, LoFreq, mpileup, Platypus, SNVer, SNVSniffer, Strelka, and VarScan) produce as output an unregularized VCF: **$root.$aligner**.**$caller**.vcf. This file is then regularised using the vcfallelicprimitives module of VCFlib, producing a final VCF for evaluation:

vcfallelicprimitives **$root.$aligner.$caller**.vcf > **$root.$aligner.$caller**.regularised.vcf.

16GT

bam2snapshot -i **$ref** -b **$root.$aligner**.bam -o **$root.$aligner.$caller**

snapshotSnpcaller -i **$ref** -o **$root.$aligner.$caller**

perl txt2vcf.pl **$root.$aligner.$caller**.txt **$root.$aligner.$caller $ref** > **$root.$aligner.$caller**.vcf

rm **$root.$aligner.$caller**.alignmentQC.txt **$root.$aligner.$caller**.snapshot **$root.$aligner.$caller**.txt

Freebayes

freebayes -f **$ref** --ploidy 1 **$root.$aligner**.bam > **$root.$aligner.$caller**.vcf

GATK

gatk HaplotypeCaller -R **$ref** -I **$root.$aligner**.bam -O **$root.$aligner.$caller**.vcf

rm **$root.$aligner.$caller**.vcf.idx

LoFreq

lofreq call -f **$ref** -o **$root.$aligner.$caller**.vcf **$root.$aligner**.bam

mpileup

bcftools mpileup -Ou -f **$ref $root.$aligner**.bam | bcftools call --threads 40 --ploidy 1 -mv -Ov -o **$root.$aligner.$caller**.vcf

Platypus

python Python.py callVariants --bamFiles=**$root.$aligner**.bam --logFileName=**$root.$aligner.$caller**.log --refFile=**$ref** --output=**$root.$aligner.$caller**.vcf

rm **$root.$aligner.$caller**.log

SNVer

java -jar SNVerIndividual.jar -i **$root.$aligner**.bam -r **$ref** -o **$root.$aligner.$caller**

mv **$root.$aligner.$caller**.filter.vcf **$root.$aligner.$caller**.vcf

rm **$root.$aligner.$caller**.failed.log **$root.$aligner.$caller**.indel.filter.vcf **$root.$aligner.$caller**.indel.raw.vcf **$root.$aligner.$caller**.filter.vcf **$root.$aligner.$caller**.raw.vcf

SNVSniffer

samtools view -H **$root.$aligner**.bam > **$root.$aligner**.header.sam

SNVSniffer snp -f 2 -g **$ref** -o **$root.$aligner.$caller**.vcf **$root.$aligner**.header.sam **$root.$aligner**.bam

rm **$root.$aligner**.header.sam

Strelka

python configureStrelkaGermlineWorkflow.py --bam **$root.$aligner**.bam --referenceFasta **$ref** --runDir **$root.$aligner.$caller**

python **$root.$aligner.$caller**/runWorkflow.py -m local -j 40

gunzip **$root.$aligner.$caller**/results/variants/genome.S1.vcf.gz

mv **$root.$aligner.$caller**/results/variants/genome.S1.vcf **$root.$aligner.$caller**.vcf

rm -r **$root.$aligner.$caller**

VarScan

samtools mpileup -B -q 1 -f **$ref** **$root.$aligner**.bam > **$root.$aligner.$caller**.mpileup

java -jar VarScan.v2.3.9.jar mpileup2snp **$root.$aligner.$caller**.mpileup --output-vcf 1 > **$root.$aligner.$caller**.vcf

rm **$root.$aligner.$caller**.mpileup

(2) **Comparing pipeline performance when simulating both error-free and error-containing reads**

When initially evaluating each pipeline, all reads were simulated error-free. This was in order to exclude biases at other points in the workflow, such as in DNA library preparation. To compare pipeline performance between sets of error-free and error-containing reads, a parallel set of equivalent simulations – of 3 sets of 150bp and 3 sets of 300bp paired-end reads, each at 50x base-level coverage and aligned both to the same genome from which they were simulated and to a divergent genome – were performed only for the set of *E. coli* strains (*E. coli* was chosen as it was among the most diverse of the 10 species in this study, with the greatest range of genome sizes; see Supplementary Table 5).

Error-containing reads were simulated using dwgsim v0.1.11 (https://github.com/nh13/DWGSIM) with parameters -e 0.001-0.01 (non-uniform per-base error rate increasing across the read from 0.01 to 0.1%, approximating a generic Illumina error profile) and -y 0.01 (1% probability of simulating a random DNA read). Parallel sets of error-free reads were simulated with dwgsim parameters -e 0-0 and -y 0.

dwgsim does not output the otherwise randomly generated seed used for each simulation, although does allow seeds to be provided. To ensure results were reproducible, the same seeds were provided to dwgsim as were generated during the initial set of error-free (wgsim) simulations (i.e., as given in Supplementary Table 3).

This dataset contains 4 sets of 7134 VCFs, 2 made using dwgsim error-free reads (i.e., aligned to the same genome and aligned to the representative genome) and 2 made using error-containing reads. Each dataset of 7134 VCFs comprises 2 read lengths (150 and 300bp) * 3 replicates * 29 *E. coli* strains * 41 pipelines. All VCFs were filtered according to criteria detailed in Supplementary Table 12 (and which were originally empirically selected for use with the COMPASS pipeline; see below).

The performance statistics for each pipeline are shown in Supplementary Tables 13 (for reads aligned to the same genome from which they were simulated) and 14 (for reads aligned to the representative genome).

When aligning reads to the same genome from which they were simulated, there were near-perfect correlations between F-scores obtained using error-free and error-containing reads (Spearman’s *rho* = 0.9981, p < 10^-15^; see figure below). Sequencing error does introduce a (negligible) number of false positive calls, however. The correlation between estimates of precision (positive predictive value) is, while strong, marginally weaker than the correlation between estimates of recall (sensitivity): for precision, Spearman’s *rho* = 0.9282 (p < 10^-15^) and for recall, Spearman’s *rho* = 0.9979 (p < 10^-15^). Similar results are seen when aligning reads to a divergent, representative, genome (see figure below), with equivalently strong correlations between F-score (Spearman’s *rho* = 0.9889, p < 10^-15^), precision (Spearman’s *rho* = 0.9870, p < 10^-15^) and recall (Spearman’s *rho* = 0.9862, p < 10^-15^).

These results suggest that introducing sequencing error into the simulated datasets has a negligible effect on the overall performance rank of each pipeline, even in the absence of pre-processing quality control (such as by, e.g., the read trimmer Trimmomatic [1]) which should in principle diminish error further.

This can likely be attributed to the VCF filter criteria (which are broadly similar to those recommended by a previous study for maximising SNP validation rate [2]) and/or because many of the aligners already apply internal mechanisms for accommodating error (such as ‘soft-clipping’: omitting unaligned, i.e. error-prone, terminal regions from reads and using only partial alignments between the reads and reference [3]).

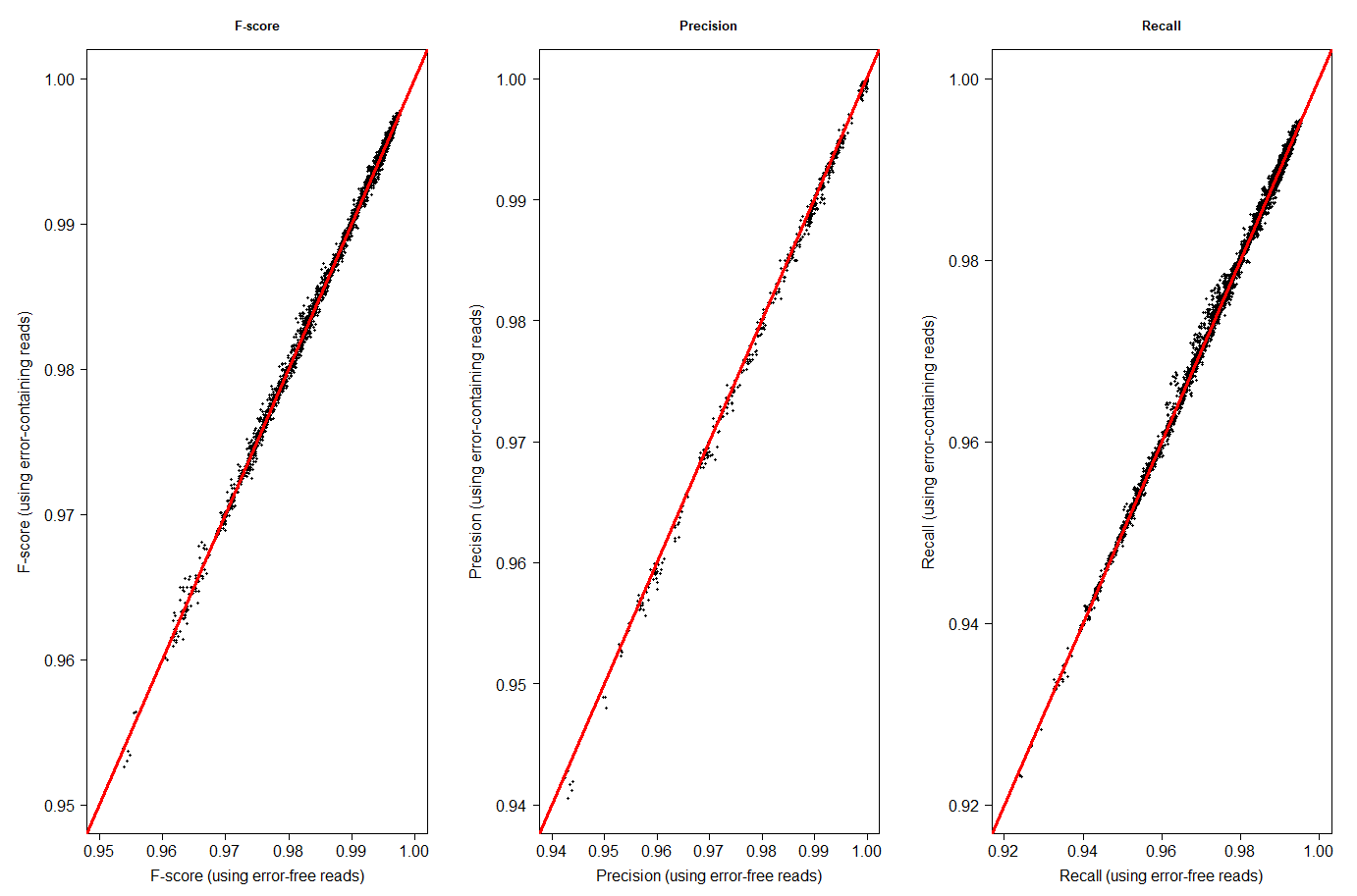

**Head-to-head performance comparison of pipelines evaluated using both error-free and error-containing reads, aligning reads to the same genome from which they were simulated.**

This figure directly compares the performance of pipelines using both error-free and error-containing reads (which have a non-uniform per-base error rate increasing across the read from 0.01 to 0.1%). In both cases, reads are simulated from 29 *E. coli* strains (detailed in Supplementary Table 2) and aligned back to the same genome. Each point represents a simulation (n = 7134, i.e. 2 read lengths [150 and 300bp] * 3 replicates * 29 *E. coli* strains * 41 pipelines). Summary statistics for each simulation are shown in Supplementary Table 13. The line y = x is shown in red.

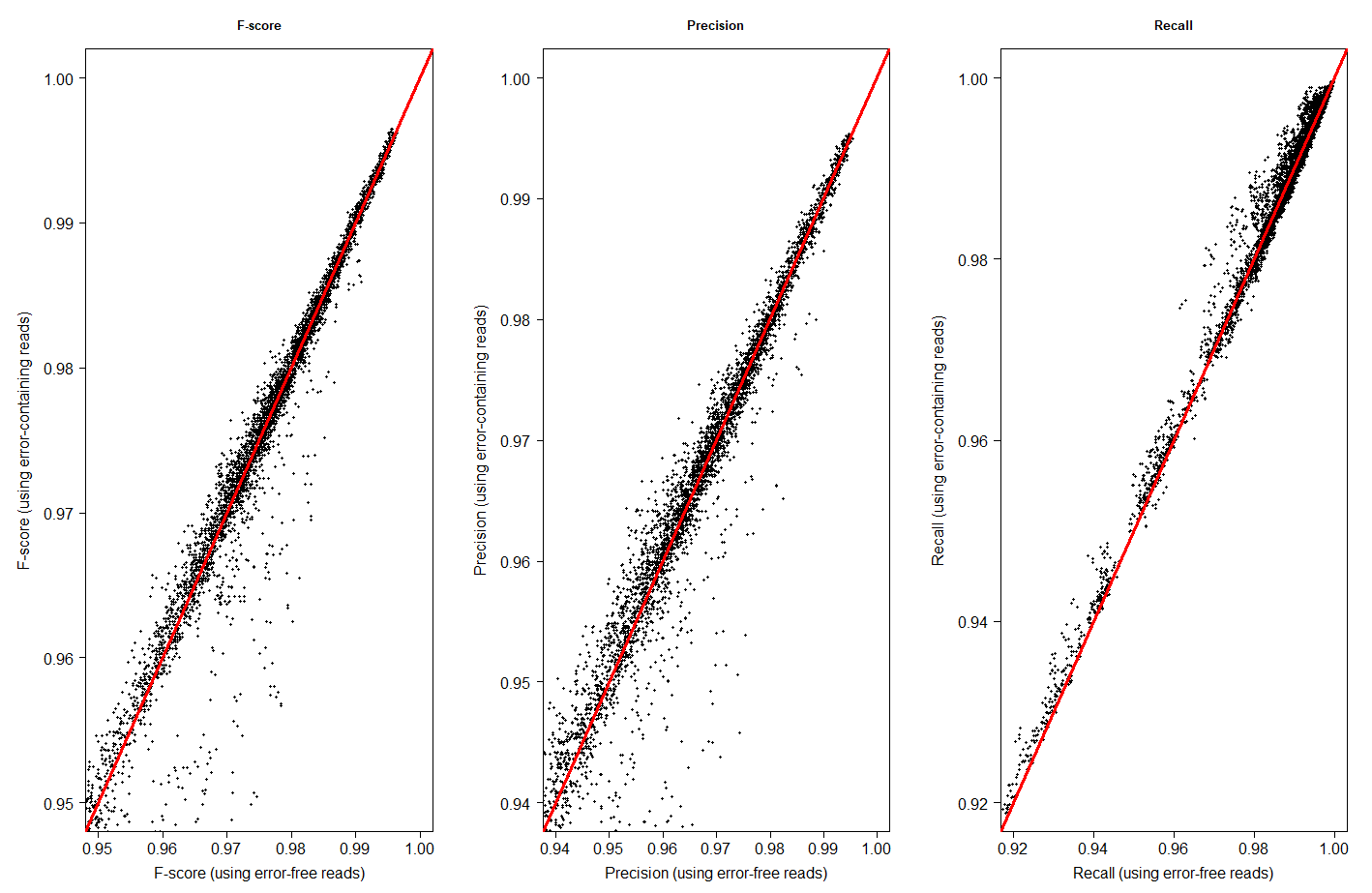

**Head-to-head performance comparison of pipelines evaluated using both error-free and error-containing reads, aligning reads to a different genome from which they were simulated.**

This figure directly compares the performance of pipelines using both error-free and error-containing reads (which have a non-uniform per-base error rate increasing across the read from 0.01 to 0.1%). In both cases, reads are simulated from 29 *E. coli* strains (detailed in Supplementary Table 2) and aligned back to a different, representative, genome. Each point represents a simulation (n = 7134, i.e. 2 read lengths [150 and 300bp] * 3 replicates * 29 *E. coli* strains * 41 pipelines). Summary statistics for each simulation are shown in Supplementary Table 14. The line y = x is shown in red.

(3) **Comparing pipeline performance with error-free reads if adding an additional BAM post-processing step of local indel realignment**

A formerly commonplace post-processing step for many aligner/caller pipelines was to locally realign around indels in the BAM, correcting for mismatching bases that could otherwise be mistaken for SNPs (these can be introduced by indels in the genome that are absent in the reference).

However, indel realignment, using the GATK modules RealignerTargetCreator and IndelRealigner, is not necessarily required for variant discovery if using a variant caller that already incorporates a haplotype assembly step, either by local *de novo* re-assembly (such as Platypus or GATK HaplotypeCaller, the successor to UnifiedGenotyper) or by building the haplotype directly from the reads (such as Freebayes). A previous study has also demonstrated that local indel realignment, even in divergent regions, had minimal impact on SNP calling accuracy [4]. To that end, the RealignerTargetCreator and IndelRealigner modules were removed from GATK v4 (and in any case are unable to re-align reads around insertions > 30bp), although this does not preclude potential added-value benefit if applied to poor-quality data (https://software.broadinstitute.org/gatk/blog?id=7847, accessed 2nd April 2019).

Consequently, we did not incorporate a routine local realignment step in each pipeline (between Picard MarkDuplicates and BuildBamIndex; see ‘second step of each pipeline’, above). To test whether local indel realignment had a quantifiable effect upon pipeline performance, we created a parallel set of simulations – of 3 sets of error-free 150bp and 3 sets of error-free 300bp paired-end reads, each at 50x base-level coverage and aligned to a divergent genome – for the diverse set of *E. coli* strains (as above). These can be directly contrasted with the error-free *E. coli* simulations in Supplementary Table 6.

To perform indel realignment, we used a deprecated version of GATK, v3.8.1 (https://github.com/broadgsa/gatk/releases, accessed 30^th^ May 2019), the last available with this functionality.

The performance statistics for each pipeline, both with- and without local indel realignment, are shown in Supplementary Tables 15. There were near-perfect correlations between F-scores irrespective of whether local indel realignment was performed (Spearman’s *rho* = 0.9993, p < 10^-15^; see figure below). Equivalently strong correlations were found between precision (Spearman’s *rho* = 0.9987, p < 10^-15^) and recall (Spearman’s *rho* = 0.9993, p < 10^-15^). While local indel realignment reduces false positive calls, so increasing precision, among the poorer-performing pipelines, the effect size of this difference is negligible (Cliff’s delta = -0.008).

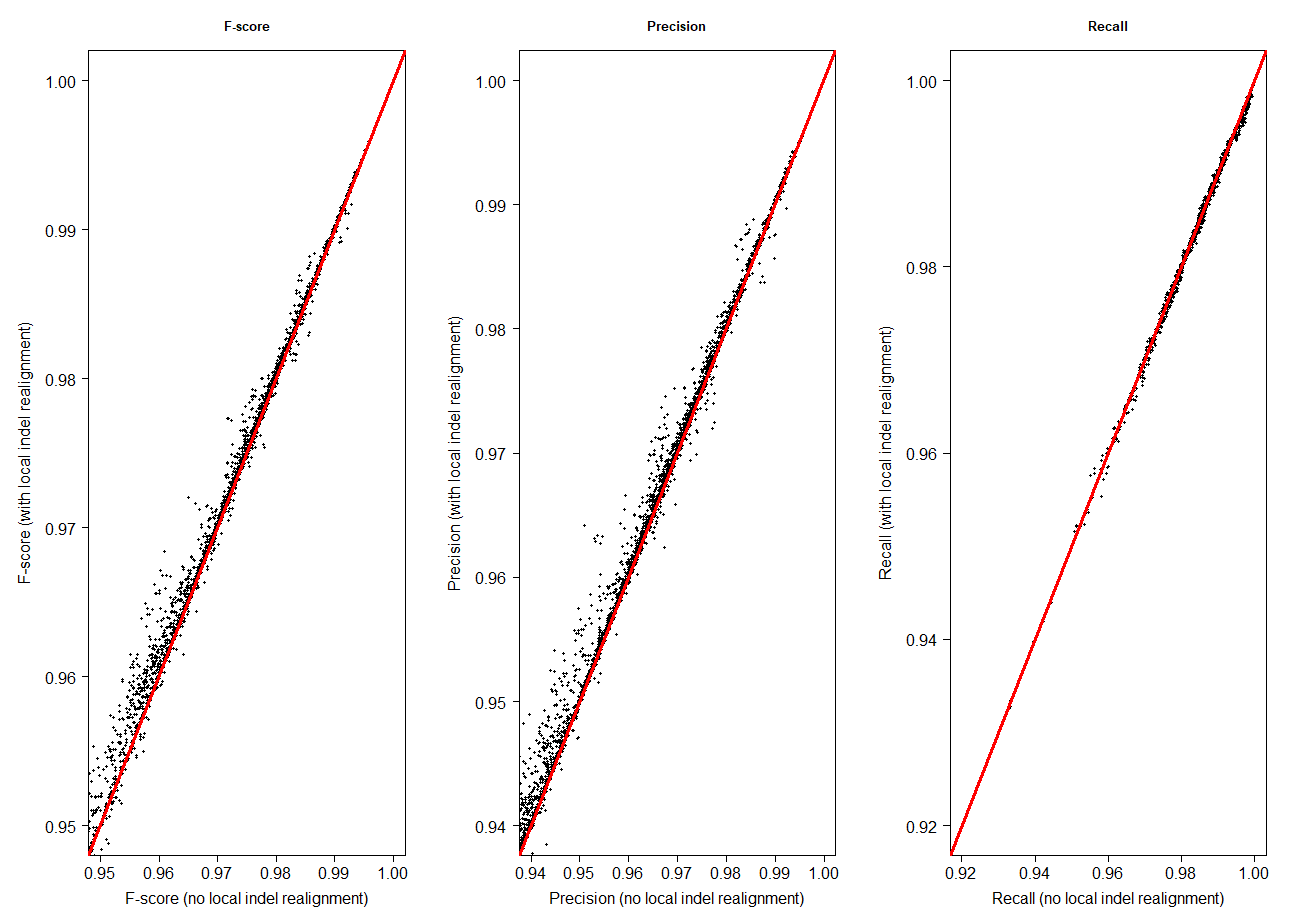

**Head-to-head performance comparison of pipelines evaluated both with and without a local indel realignment step, aligning reads to a different genome from which they were simulated.**

This figure directly compares the performance of pipelines evaluated both with and without a BAM post-processing step of local indel realignment. In both cases, reads are simulated from 29 *E. coli* strains (detailed in Supplementary Table 2) and aligned back to a different, representative, genome. Each point represents a simulation (n = 7134, i.e. 2 read lengths [150 and 300bp] * 3 replicates * 29 *E. coli* strains * 41 pipelines). Summary statistics for each simulation are shown in Supplementary Table 15. The line y = x is shown in red.

(4) **Programs not evaluated in this study**

As there are comparatively few variant calling evaluations using bacterial data, we sought to test a broad range of programs, specifically those that are in common use, under active development and which produce comparable output in standard formats (BAM, VCF).

To create a shortlist of candidate aligners and variant callers, we used similar criteria to a previous study [5], excluding from consideration programs that were formally deprecated (such as the aligner SHRiMP2 [6] and caller Isaac [7]), not being actively developed (having no meaningful version update in the last 2-3 years), not intended for use with Illumina reads (such as drFAST [8] and lordFAST [9], which are intended only for use with AB SOLiD colour space and PacBio reads, respectively), only capable of calling specific types of variant (such as the indel-only Dindel [10], SNP-only SNPSVM [11] and structural variant-only BreakDancer [12]), having hardware requirements which could not be met (such as SOAP3-dp [13], which requires a CUDA-enabled GPU), requiring matched samples as input (a common requirement of many somatic variant callers [14]), or which made assumptions of the input which were not met by the simulated datasets (such as DiscoSnp [15] and DiscoSnp++ [16], which are intended to call only isolated SNPs, i.e. those distant both up- and downstream from any other polymorphism by at least *k* nucleotides). We also excluded DeepVariant [17], a TensorFlow machine-learning based variant caller, as this required a training model made using a deep neural network. However, there is currently no production-grade DeepVariant training pipeline (the default training model supplied with DeepVariant is based on human data), nor are there high-confidence, non-simulated, bacterial truth sets on which to train it.

Self-contained variant calling pipelines, such as Snippy (https://github.com/tseemann/snippy), may incorporate programs beyond an aligner and caller alone, use the same aligners and callers independently included in this study and/or employ their own internal quality-checking mechanisms. For the purpose of this study – which evaluates SNP calling when the reference genome is not explicitly known – we sought to use all aligners and callers uniformly, with equivalent quality-control steps applied to all reads. However, although it would minimise an additional source of variation to exclude self-contained pipelines, such as Snippy, from the evaluation, it would not be a realistic use case – this is because it is increasingly common informatics practice to use composite, or containerised, approaches to particular tasks. (Note that although the Breseq [18] pipeline is another such approach, employing the aligner Bowtie2 [19] and calling variants from read pileups, it was excluded from consideration because it outputs results in Breseq-specific ‘GenomeDiff’ format rather than as a VCF).

To that end, while direct comparisons of any aligner/caller pipeline with self-contained pipelines are possible, the results must be interpreted with caution. This is because it is in principle possible to improve the performance of the former through additional quality control steps (that is, it is not necessarily the aligner or caller alone to which any relative reduction in performance may be attributed).

We aimed for this evaluation to be extensive but it is neither realistic nor computationally tractable to be exhaustive. For practicality, we excluded numerous short read aligners including but not limited to Bowtie [20], Bowtie2 [19], Cushaw3 [21], GEM [22], GSNAP [23], HISAT2 [24], Masai [25], MOSAIK [26], mrFAST [27], mrsFAST [28], mrsFAST-Ultra [29], NextGenMap [30], RazerS [31], RazerS3 [32], SeqAlto [33], SMALT (http://www.sanger.ac.uk/science/tools/smalt-0), SRmapper [34], and ZOOM [35].

Although it is not possible to benchmark all (or even a majority) of tools on all possible metrics, one of the core findings of this evaluation is that the Mash distance between the reads and the reference genome is especially critical to pipeline performance. To that end, there is no compelling reason to believe that those tools which have not been evaluated – assuming they were not designed for this scenario – will differ greatly in performance from those that have.

(5) **Overview of the COMPASS (Complete Pathogen Sequencing Solution) pipeline and associated VCF parsing criteria**

The filter criteria used to parse each VCF (detailed in Supplementary Table 12) were adapted from those empirically developed for use in the COMPASS pipeline. COMPASS development began in 2010 by the Modernising Medical Microbiology (MMM) team (http://modmedmicro.nsms.ox.ac.uk/), with the aim to process a high volume of sequencing data into a form suitable for clinical reports. The rationale behind the development of COMPASS was that although a variety of tools were available for individual sequence analyses (for reviews, see [36, 37]), an end-to-end software solution, for use in routine clinical practice, was not. COMPASS was developed for this purpose on a cloud-based platform to offer the capability of future growth.

While not independently published, results generated by COMPASS have been utilised in numerous publications, demonstrating the strength and applicability of the pipeline [38-46]. COMPASS comprises several processes that download raw sequencing data for a given sample, undertake quality control to ensure the sequencing has worked, assembles a reference genome, maps reads to this reference, and then calls variants from the mapped read data. To do so, COMPASS employs BLAST+ v 2.2.23 [47], Bowtie2 v2.2.23 [19], BWA v0.7.5a [48], FastQC v0.11.2, GATK v1.4.21 [49], Kraken v0.10.06 [50], Mykrobe Predictor v0.3.1-0-g87 [51], Picard Tools v1.31, SAMtools v0.1.19 [52], and Stampy v1.0.23 [53], with bespoke linking scripts written in Python v2.7.1 and using standard Python libraries as required.

Using per-site statistics from the subsequent VCF and empirically-determined filter criteria, COMPASS then re-calls each base in the assembled sample with respect to the reference genome, outputting an expanded VCF that contains calls for every site. As such, by extracting the ‘alt’ column of this VCF, a FASTA file can be obtained for each sample. COMPASS attempts base calling only for sample sites that have homologues in the reference genome, so the expanded VCF (and subsequent FASTA) always contains the same number of sites as the reference genome. Should one of the COMPASS filter criteria not be met, an ‘N’ call is made instead, indicating insufficient, inconclusive or conflicting information for that site.

The set of filter criteria employed by COMPASS underwent several revisions, of which only the final set of criteria for the current version are used in this evaluation. Consistency in variant calling between different sets of criteria was assessed by resequencing isolates from the same bacterial colonies on different flow cells as technical replicates. The criteria used for evaluation were the number of discordant calls between each pair of replicates, which measured how often a variant was incorrectly detected between identical genomes (i.e. false positive rate), and the proportion of the genome called between a pair of replicate sequences. This indirectly measured the “false negative” rate – assuming no-calls occur randomly, the call rate would be proportional to how often a true variant was not detected between a pair of genomes.

Briefly, COMPASS requires all variant calls to be homozygous under a diploid model. In all versions of COMPASS, variant calls were made only in non-repetitive regions of the core genome, with repetitive regions identified by self-self BLAST and subsequently masked. For the purpose of this evaluation, we did not exclude calls within these regions as this would require appending VCFs prior to filtering, i.e. filtering calls on the basis of information (a ‘within a repetitive region’ flag) not initially output by any aligner/caller combination. In addition, the retention of repetitive regions, and their associated lower-confidence calls, would have a systematic negative effect on the performance of each pipeline.

The initial set of COMPASS filter criteria (as used in, for instance, [54]) required that a SNP have a consensus of 75% of the mapped reads and support from at least five reads, including one in each direction, at a site with a depth of high-quality (mapping quality Phred score > 20) coverage between the 2.5 and 97.5 percentiles of all sites for that isolate (i.e. the SNP does not occur at a site with unusual depth). Sites where minority variants represented more than 10% of read depth were also defined as ‘mixed’, with no base called. Variant calls were also not made if they occurred within 12bp of another nucleotide variant or indel. The most recent set of COMPASS filters, used in this evaluation, removed the ‘unusual depth’ and ‘nearby variant’ criteria as they were found to increase mean genome coverage without affecting estimates of genetic relatedness or introducing false positive calls [55]. The minimum requirements for coverage, Phred quality score, and call consensus are retained.
